# The domain-separation low-dimensional language network dynamics in the resting-state support the flexible functional segregation and integration during language and speech processing

**DOI:** 10.1101/2022.06.19.496753

**Authors:** Binke Yuan, Hui Xie, Zhihao Wang, Yangwen Xu, Hanqing Zhang, Jiaxuan Liu, Lifeng Chen, Chaoqun Li, Shiyao Tan, Zonghui Lin, Xin Hu, Tianyi Gu, Junfeng Lu, Dongqiang Liu, Jinsong Wu

**Affiliations:** Key Laboratory of Brain, Cognition and Education Sciences, Ministry of Education, China; Institute for Brain Research and Rehabilitation, South China Normal University, Guangzhou, China; Department of Psychology, The University of Hong Kong, Hong Kong, China; Shenzhen Key Laboratory of Affective and Social Neuroscience, Magnetic Resonance Imaging Center, Center for Brain Disorders and Cognitive Sciences, Center for Brain Disorders and Cognitive Sciences, Shenzhen University, Shenzhen 518060, China; Department of Biomedical Sciences of Cells & Systems, Section Cognitive Neuroscience, University Medical Center Groningen, University of Groningen, Groningen, The Netherlands; Center for Mind/Brain Sciences (CIMeC), University of Trento, Trento 38123, Italy; Glioma Surgery Division, Neurologic Surgery Department, Huashan Hospital, Shanghai Medical College, Fudan University, Shanghai, China; Brain Function Laboratory, Neurosurgical Institute of Fudan University, Shanghai, China; 9Shanghai Key Laboratory of Brain Function Restoration and Neural Regeneration, Shanghai, China; Research Center of Brain and Cognitive Neuroscience, Liaoning Normal University, Dalian, China; Key Laboratory of Brain and Cognitive Neuroscience, Liaoning Province, Dalian, PR China

## Abstract

Modern linguistic theories and network science propose that the language and speech processing is organized into hierarchical, segregated large-scale subnetworks, with a core of dorsal (phonological) stream and ventral (semantic) stream. The two streams are asymmetrically recruited in receptive and expressive language or speech tasks, which showed flexible functional segregation and integration. We hypothesized that the functional segregation of the two streams was supported by the underlying network segregation. A dynamic conditional correlation approach was employed to construct frame-wise time-varying language networks and investigate the temporal reoccurring patterns. We found that the time-varying language networks in the resting-state robustly clustered into four low-dimensional states, which dynamically reconfigured following a domain-separation manner. Spatially, the hub distributions of the first three states highly resembled the neurobiology of primary auditory processing and lexical-phonological processing, motor and speech production processing, and semantic processing, respectively. The fourth state was characterized by the weakest functional connectivity and subserved as a baseline state. Temporally, the first three states appeared exclusively in limited time bins (∼15%), and most of the time (> 55%), the language network kept inactive in state 4. Machine learning-based dFC-linguistics prediction analyses showed that dFCs of the four states significantly predicted individual linguistic performance. These findings suggest a domain-separation manner of language network dynamics in the resting-state, which forms a dynamic “meta-networking” (network of networks) framework.

**Highlights:**

1. The time-varying language network in the resting-state is robustly clustered into four low-dimensional states.
2. Spatially, the first three dFC states are cognitively meaningful, which highly resemble the neurobiology of primary auditory processing and lexical-phonological representation, speech production processing, and semantic processing, respectively.
3. Temporally, the first three states appeared exclusively in limited time bins (∼15%), and most of the time (> 55%), the language network kept inactive in state 4.
4. A dynamic “meta-networking” framework of language network in the resting-state is proposed.

## 1 Introduction

Linguistic theories propose that language and speech processing composes of phonological processing (Wagner and Torgesen, 1987), semantic processing (Binder et al., 2009; Collins and Loftus, 1975; Patterson et al., 2007), and sentence/syntactic processing (Matchin and Hickok, 2020). Brain regions implicated in these domain-specific representations are now thought to span laterally in the left frontal, temporal and parietal cortices (Fedorenko and Thompson-Schill, 2014; Fridriksson et al., 2016; Lu et al., 2017; Lu et al., 2021; Price, 2012; Vigneau et al., 2006; Wu et al., 2015; Zhao et al., 2021), which are connected by the subcortical white matter fiber tracts (Dick et al., 2014; Hula et al., 2020; Saur et al., 2008; Yagmurlu et al., 2016; Zhang et al., 2021). Researchers proposed that the language or speech network is organized into hierarchical, segregated large-scale subnetworks, with a core of dorsal (phonological) stream and ventral (semantic) stream (Duffau et al., 2014; Fridriksson et al., 2016; Hickok and Poeppel, 2007; Hodgson et al., 2021; Hula et al., 2020; Roelofs, 2014; Ueno et al., 2011).

Functionally, the phonological stream participates in auditory- and articulatory-phonological processing, while the semantic stream is primarily concerned with conceptual-semantic processing. Anatomically, distinct anatomical boundaries were revealed. The phonological stream is mainly distributed in the left prefrontal cortex and superior temporal gyrus, while the ventral stream is mainly distributed in bilateral temporal and parietal lobes (Fridriksson et al., 2016; Hickok and Poeppel, 2004, 2007; Hodgson et al., 2021). The implication of the two distinguishable streams is task-dependent. For auditory-to-conceptual tasks, such as speech comprehension, the ventral stream was predominantly recruited (Barch et al., 2013; Nastase et al., 2021; Sefcikova et al., 2022). The speech signal was hierarchically processed, involving lexical-phonological representation, lexical/syntactic representation, and semantic representation (Binder, 2017; Hickok, 2022). While, for tasks including both conceptual-to-auditory and auditory-to-motor, such as speech production, both the dorsal and ventral streams were recruited (Hassan et al., 2015; Lu et al., 2021; Price, 2012). Semantic or conceptual preparation and lexical selection were firstly processed in the ventral stream, and then lexical-phonological retrieval, morphological and phonemic sequences processing, and articulation were successively processed in the dorsal motor and speech production network (Duffau et al., 2014; Hickok, 2022; Hu et al., 2021). Generally, receptive language only involves a subset of the language network (mainly the ventral stream) (Dronkers et al., 2004), while expressive language hierarchically involves the whole language network. Thus, the two streams are more likely to form a hierarchical parallel architecture (Matchin et al., 2022). The asymmetrical recruitment also well explained the double dissociations of expressive and receptive language deficits (Fridriksson et al., 2018; Matchin et al., 2022). For example, damage to the dorsal stream resulted in reception-spare syndromes, such as apraxia of speech (Kent and Rosenbek, 1983; Ogar et al., 2005), Broca’s aphasia (Mohr et al., 1978), and conduction aphasia (Benson et al., 1973; Bernal and Ardila, 2009). However, damage to the ventral stream resulted in both expressive and receptive deficits, such as Wernicke’s aphasia (Robson et al., 2019; Weiller et al., 1995) and word deafness (Buchman et al., 1986; Poeppel, 2001). All these observations suggested that the two streams can flexibly and dynamically segregate or integrate for diverse cognitive processes.

From a network perspective, functional segregation and integration were supported by the underlying network segregation and integration (He and Evans, 2010; Park and Friston, 2013; Sporns, 2013; Yuan et al., 2017). The brain is an inherently dynamic system to spatially and temporally integrates distributed but relatively specialized networks (Avena-Koenigsberger et al., 2017; Cabral et al., 2017; Herbet and Duffau, 2020; Park and Friston, 2013; Pezzulo et al., 2021). Functional network reconfiguration occurs in multi-temporal and topological scales and during both rest and task states (Betzel and Bassett, 2017; Hutchison et al., 2013). The dynamic nature endows the brain flexibly and dynamically segregates or integrates during diverse cognitive processes (Cohen, 2018; Medaglia et al., 2015). Motivated by the highly dynamic and segregated phenomena of spontaneous brain activity (Avena-Koenigsberger et al., 2017; Cabral et al., 2017; Cohen and D’Esposito, 2016; Hearne et al., 2017; Ma and Zhang, 2018; Pezzulo et al., 2021), in task-free condition, we hypothesized that the moment-to-moment language network dynamics would be functionally segregated, which was essential for flexibly and dynamically integrated during language and speech processing. The moment-to-moment language network dynamics can be clustered into several temporal reoccurring states (Hutchison et al., 2013), which reflect the meta-networking organization of language network (Herbet and Duffau, 2020). To this aim, we employed a dynamic conditional correlation approach (Lindquist et al., 2014) to construct the frame-wise dynamic network and investigate the spatiotemporal characteristics of language network dynamics in the resting-state. Moreover, we also hypothesized that the meta states of language network would be cognitive- or behavioral relevance, which could be (partly) predictive for off-line linguistic performance (Eichenbaum et al., 2021; Pezzulo et al., 2021).

## 2 Materials and Methods

### 2.1 fMRI datasets

#### Discovery dataset

The dataset was part of the 1000 Functional Connectomes Project and is available at http://fcon_1000.projects.nitrc.org. Six subjects were excluded due to preprocessing errors and the remaining 192 subjects (118 females; age, 18–26 years old, mean ± STD = 21.17 ± 1.83) were analyzed. The scanning parameter details can be found at http://fcon_1000.projects.nitrc.org/fcpClassic/FcpTable.html. Participants were instructed to keep their heads still, and close their eyes but be awake.

#### Validation dataset

To ensure robustness, we adopted a large dataset from the Human Connectome Project (HCP) 900 Subject Release (see Van Essen et al., 2012 for dataset details). The first run data with phase encoding in a right-to-left direction were used. The inclusion criteria and data preprocessing have been detailed in Wang et al. (2021). Finally, 501 participants (276 females; age, mean ± STD = 28.51 ± 3.62) were analyzed. The HCP dataset includes a battery of behavioral/cognitive tests (Barch et al., 2013). Two language-related tests (Oral Reading Recognition Test [ORRT] and Picture Vocabulary Test [PVT]) were adopted to investigate the behavioral relevance of the language network dynamics (section 2.7).

The corresponding Ethics Committee approved the study. All procedures followed the Declaration of Helsinki. Written informed consent was obtained from each participant.

### 2.2 Data preprocessing

Discovery dataset: 1) The first 10 volumes were discarded; 2) slice-timing; 3) head motion correction. No participant exhibited head motion > 3 mm maximum translation or 3° maximum rotation; 4) The motion-corrected functional images were co-registered to the 3D-T1 images and then spatially normalized into MNI space by applying the deformation field as estimated by segmentation; 5) The normalized images were spatially smoothed using a Gaussian kernel (full-width-at-half-maximum = 6 mm); 6) detrend; 7) nuisance signals regression. 36 parameters, including the six parameters of translations and rotations + white matter/cerebrospinal fluid/global mean time courses (9 parameters), plus their temporal derivative (9 parameters) and the quadratic term of 18 parameters (Ciric et al., 2017; Satterthwaite et al., 2013)— were removed by linear regression from each voxel’s time course (https://github.com/sandywang/RegressOutForRfMRIwithTumor); 8) temporal band-pass filtering (0.01–0.1 Hz).

#### Validation dataset

The preprocessing steps were the same as Wang et al.(2021), which are summarized in the Supplementary materials.

### 2.3 Network definition of language processing

The putative network for language and speech processing was defined in the Human Brainnetome Atlas (Fan et al., 2016). The atlas included 246 regions parcellated based on white matter structural connectivity. For each parcel, Fan and colleagues performed meta-analyses according to the behavioral domain and paradigm-class meta-data labels from the BrainMap database. We extracted all parcels that exhibited significant activation in language- or speech-related behavioral domains or paradigm classes (e.g., semantic, word generation, reading, or comprehension). Most of these cortical parcels are in the left hemisphere and their extents are highly similar to those identified by other researchers (Fedorenko et al., 2010; Lombardo et al., 2018; Rampinini et al., 2017; Vigneau et al., 2006). Considering bilateral language processing (Chai et al., 2016; Hodgson et al., 2021; Lerner et al., 2011; Muller and Meyer, 2014; Rice et al., 2015; Siegel et al., 2016) and the recruitment of right hemisphere language regions for compensation in patients, parcels of homologs in the right hemisphere were also included. Altogether, a symmetric language network atlas (Figure 1) including 68 cortical ROIs (34 ROIs in each hemisphere) was defined (Yuan et al., 2022), including the superior, middle, and inferior frontal gyrus (SFG, MFG, and IFG, respectively), the ventral parts of the precentral gyrus (PrG) and postcentral gyrus (PoG) (Lu et al., 2021), the superior, middle and inferior temporal gyrus (STG, MTG, and ITG, respectively), the posterior superior temporal sulcus (pSTS), inferior parietal lobule (IPL), the fusiform gyrus (FuG), and the parahippocampus gyrus (PhG). The coordinates (in Montreal Neurologic Institute space) and meta results of each parcel are summarized in Supplementary Table 1.

**Figure 1.**
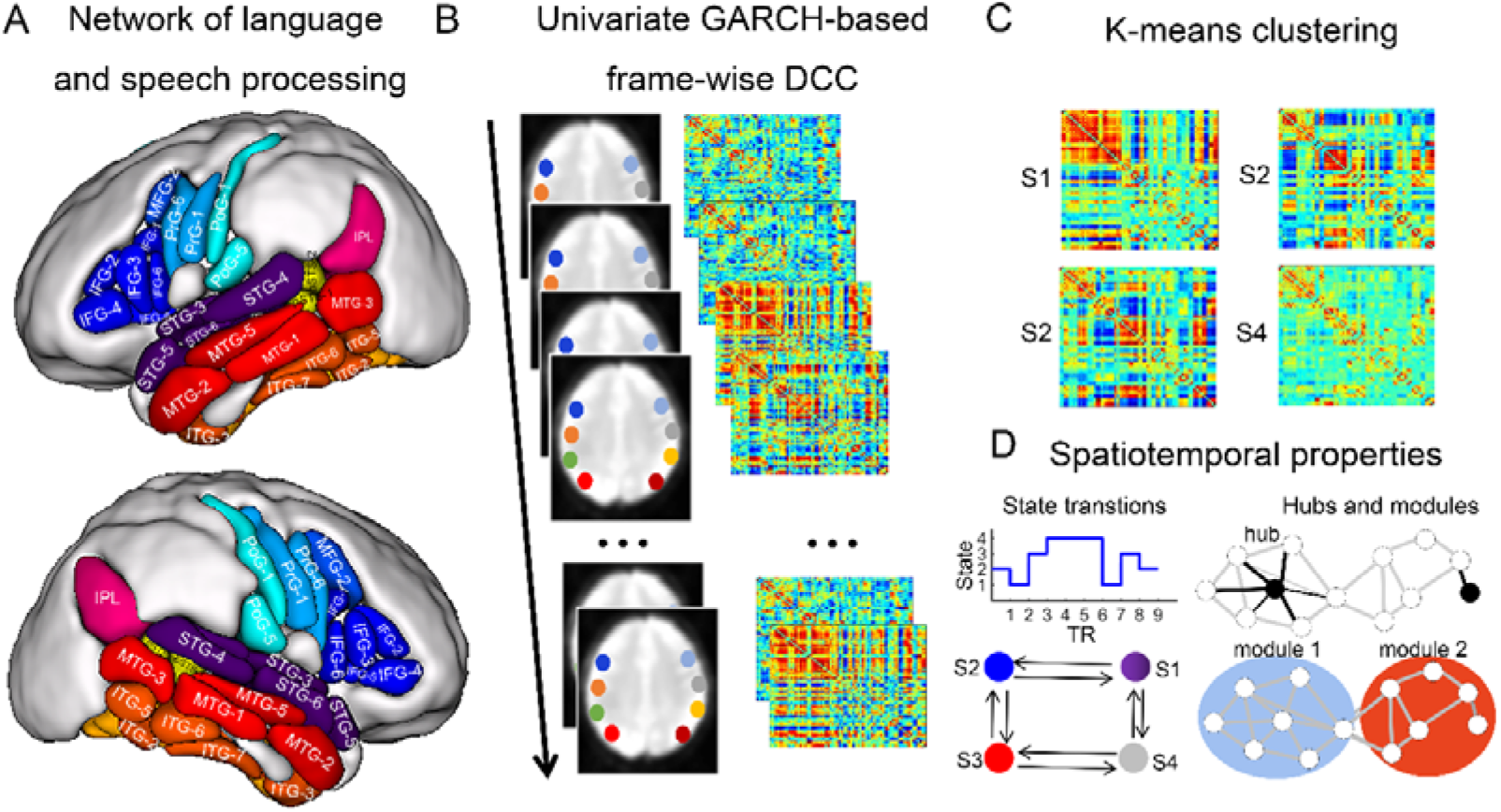
Schematic depicting the frame-wise language network dynamics analyses. A: A symmetric language network atlas (68 nodes, 34 in each hemisphere) was defined on the Human Brainnetome Atlas (Fan et al., 2016) according to the behavioral domain and paradigm-class meta-data labels from the BrainMap database (http://brainmap.org/). B: Univariate GARCH-based (Generalized AutoRegressive Conditional Heteroskedasticity) dynamic conditional correlation algorithm was adopted to construct the frame-wise dynamic networks of each subject. C: The dynamic language networks from all subjects were clustered using the k-means algorithm, yielding cluster centroids and cluster membership assignment for all frames. D: The spatiotemporal properties of the temporal reoccurring states, including the nodal degree centrality, network module, overall state frequency, and between-state transition probabilities were calculated.

### 2.4 Frame-wise time-varying language network construction using DCC

To identify the temporal reoccurring states of the language network, dynamic conditional correlation (DCC) approach was adopted to construct the frame-wise time-varying language network (Figure 1) (Lindquist et al., 2014) (https://github.com/caWApab/Lindquist_Dynamic_Correlation/tree/master/DCC_toolbox). It has been shown that DCC outperformed the sliding-window approach in tracking dynamic correlations (Choe et al., 2017; Lindquist et al., 2014).

DCC is a variant of the multivariate GARCH (generalized autoregressive conditional heteroscedastic) model (Engle, 1982; Lebo and Box-Steffensmeier, 2008), which has been shown to be particularly effective for estimating both time varying variances and correlations. All the parameters of DCC are estimated through quasi-maximum likelihood methods and require no ad hoc parameter settings. Next, we firstly introduce the concept of conditional covariance matrix (Eqs. (1) – (7)), and then we introduced how did the DCC algorithm estimate the conditional covariance matrix (Eqs. (7) – (14)).

#### Conditional covariance matrix

Let 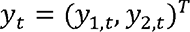, while *y*_1,*t*_ and *y*_2,*t*_ are two time series we are interested.

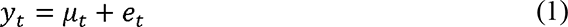

where *µ_t_* = (*µ*_1,*t*_, *µ*_2,*t*_)*^T^*, is the conditional mean of *y_t_* using all information in the time series observed up to time *t*, denoted *E_t-1_*(*y_t_*). *e_t_* is the noise term and its conditional covariance matrix at time *t* can be expressed as:

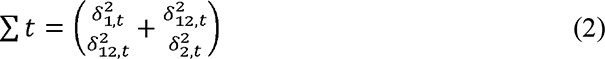

Here, the diagonal terms represent the conditional variance of *y_i,t_* using all information in the time course observed up to time 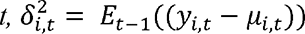 for *i =* 1, 2. The off-diagonal terms represent the conditional correlation coefficient,

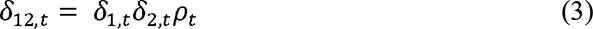

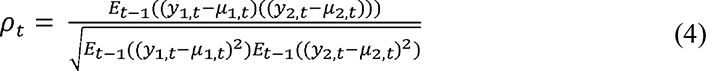

The conditional correlation at time *t* relies on information that is observed up to time *t* − 1.

If *µ_t_* = 0 and *y_t_* = *e_t_*, then *ρ_t_* is:

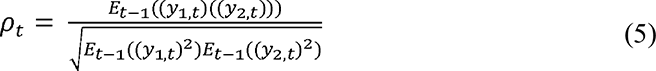

The conditional covariance matrix defined in Eq. (2) can alternatively be written in matrix form as

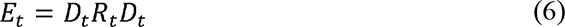

where *D_t_* is a diagonal matrix consisting of the conditional standard deviations of the time series, i.e., 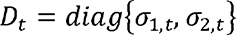 and *R_t_* is the correlation matrix, 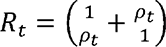. Thus,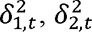, and *ρ_t_* are the three components that need to be estimated in the conditional covariance matrix.

The DCC algorithm consists of two steps. For two univariate time series, the standardized residuals are firstly computed by using univariate GARCH (1,1) models (Eqs. (7) and (8)). Based on the standardized residuals (Eqs. (9) – (11)), an EWMA-type (exponential weighted moving average) method (Eqs. (12) – (14)) is applied to compute a non-normalized version of the framewise correlation matrix (i.e., the dynamic functional connectivity matrix, dFC).

#### Univariate GARCH (1,1) model

GARCH processes are often used to model volatility in univariate time series (Engle, 1982). GARCH models express the conditional variance of a single time series at time *t* as a linear combination of the past values of the conditional variance and of the squared process itself. Let *y_t_* is a univariate process:

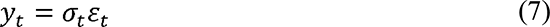

where *ε_t_* is a *N* (0, 1) random variable and *σ_t_* represents the time-varyingvariance term we seek to model. In a GARCH (1,1) process the conditional variance is expressed as

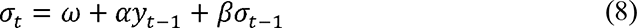

where 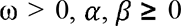 and *α* + *β* < 1. Here, the term *α* controls the impact of past values of the time series on the variance and *β* controls the impact of past values of the conditional variance on its present value.

The parameters of the GARCH model can be estimated using DCC model. Let *y_t_* = *∈_t_* is a bivariate mean zero time series with conditional covariance matrix Σ*_t_*.

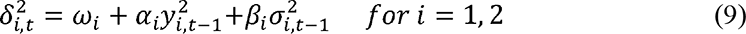

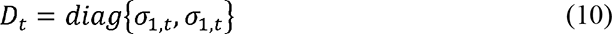

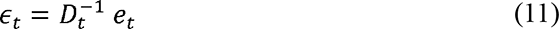

Where *∈_t_* are the standardized residuals.

#### Exponential Weighted Moving Average (EWMA)

The EWMA approach applies declining weights to the past observations in the time series based on a parameter *λ*, and is based upon the following recursion:

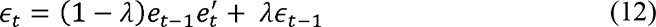

This approach places the most weight on recent observations, and for each step away from *t* values becomes gradually down-weighted by a factor *λ* (between 0 and 1), before eventually being removed from further computations. The *λ* can be estimated through maximum likelihood estimation.

Based on the standardized residuals, a non-normalized version of the framewise correlation matrix *R_t_* was estimated:

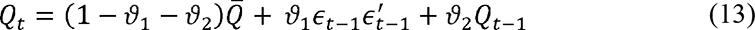

where 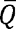 represents the unconditional covariance matrix of *∈_t_* and 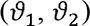 are non-negative scalars satisfying 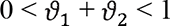.

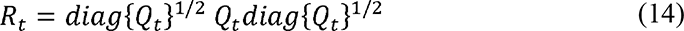

K-means clustering was adopted to decompose the dFC matrices into several low-dimensional and temporal-reoccurring connectivity states. The optimal number of clusters *k* was estimated based on the elbow criterion, i.e., the ratio between the within-cluster distance to between-cluster distances (Allen et al., 2014). L1 distance function (‘Manhattan distance’) was implemented to assess the point-to-centroid distance. Each frame was finally assigned to one of these connectivity states.

To assess the similarity and correspondence of network states between datasets, we adopted the Munkres assignment algorithm (Munkres, 1957) and calculated the spatial correlation coefficient.

### 2.5 Spatiotemporal properties of dFC states

To investigate the spatial properties of dFC states, we calculated the following metrics: 1) Nodal degree. The centroid for each state was taken as a graph and the nodal degree for a given node was calculated by summing up the correlation coefficients of all the suprathreshold connections that connected with this node. A correlation threshold, *r* > 0.2, was used to eliminate weak correlations possibly arising from noise. A node was identified as a hub if its nodal functional connectivity strength was higher than the mean nodal strength of all the identified states (Figure 1). We also assessed the robustness of our results by adopting another two correlation thresholds, *r* > 0.1 and *r* > 0.3. 2) Modularity. For each state, modularity analysis was performed on the thresholded (*r* > 0.2) binary matrix using a spectral optimization algorithm (Newman, 2006) (Figure 1). The significance of modularity of the functional brain networks was assessed by comparing it with that of 100 node-and degree-matched random networks.

To investigate the temporal properties of dFC states, we calculated the following metrics: 1) Overall state frequency, i.e., the percentage of a state occurring in all frames. We concatenated all subjects’ window membership vectors and computed the percentage of the total numbers of a given state to the total numbers of windows. Note that the overall state frequency is proportional to the total dwell times of a state in all subjects. 2) Between-state transition probability, i.e, whether a state transition occurs at time *t* + 1, given that the current state is *i*. The between-state transition probabilities were encoded in a transition matrix (Figure 1). For each subject, we first counted the numbers of transitions between each pair of states (e.g., from state *i* to state *j*), and then normalized these values by dividing the total number of state *j* (the column sum equals to 1).

As a complementary analysis, we also calculated the static functional connectivity (sFC) matrix using the Pearson correlation coefficient. We then calculated the nodal functional connectivity strength and modular assignment.

### 2.6 Frame-wise time-varying whole-brain dynamics using DCC

To investigate whether the observed language dynamics were parts of or primarily driven by the whole brain dynamics, we performed whole-brain dFC analyses using the Human Brainnetome Atlas (n = 246). Considering the computation cost, the whole-brain analysis was only performed on a subset of subjects (n = 61) from the Discovery dataset.

### 2.7 Machine learning-based dFC-linguistics prediction model

To investigate the behavioral relevance of dFCs, we investigated the relationship between dFCs and linguistics performances by constructing multivariate machine learning-based dFC-linguistics prediction models. The relevance vector regression (RVR) algorithm and linear kernel function were adopted (Cui and Gong, 2018; Yuan et al., 2019). RVR has no algorithm-specific parameter and thus does not require extra computational resources to estimate the optimal algorithm-specific parameters (Tipping, 2000).

Two language-related tests (Oral Reading Recognition Test [ORRT] and Picture Vocabulary Test [PVT]) from the HCP dataset (Barch et al., 2013) were used as predicted labels. The task descriptions, scoring process, and interpretations of each test were described in the Supplementary Materials. We used the age-adjusted scores with a mean of 100 and a standard deviation (SD) of 15 using the NIH National Norms toolbox. The criteria for outliers is out of 3 S.D of the group mean. One subject of ORRT test and one subject of PVT test were excluded.

Leave-one-out-cross-validation (LOOCV) was used to calculate the prediction accuracy (the Pearson correlation coefficient between the predicted and actual labels). In each turn of the LOOCV, one subject was designated as the test sample and the remaining subjects were used to train the model. The predicted score was then obtained by the feature matrix of the tested sample.

The significance level was computed based on 1000 permutation tests. For each permutation test, the prediction labels were randomized, and the same RVR prediction process as used in the actual data was carried out. After 1000 permutations, a random distribution of accuracies was obtained and the *P* value was correspondingly calculated: 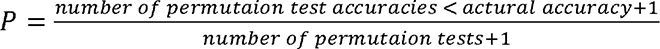. As a complementary analysis, we also constructed sFC-linguistics prediction models.

Note that an initially univariate feature-filtering step was performed to improve the model performance (Dosenbach et al., 2010). In each LOOCV fold, the correlation (Pearson’s *r*) of each of the connections with the independent variable (i.e., linguistics scores) was computed on the training set. The features were then separately ranked by the significant *p* values in ascending order. Multi p values (0.1, 0.05, 0.01) were adopted to maximize the model performance.

## 3 Results

### 3.1 The low-dimensional temporal reoccurring states of language network in the resting-state

For each subject, frame-wise DCC yielded 215 (215 volumes) functional connectivity matrices. By applying the k-means clustering algorithm and the elbow criterion to all subjects’ matrices (192 subjects * 215 frames), four temporally reoccurring dFC states with distinct patterns were identified (Figure. 2). State 1 was characterized by moderate to high positive connectivity between nodes in the bilateral PrG, PoG, and STG, but weak or moderate negative connectivity between the prefrontal nodes and temporal nodes. However, in state 2, strong positive connectivity among the prefrontal nodes was observed. State 3 distinguished itself from states 1 and 2 to strong connectivity among temporal nodes. State 4 showed an overall weak connectivity pattern.

**Figure 2.**
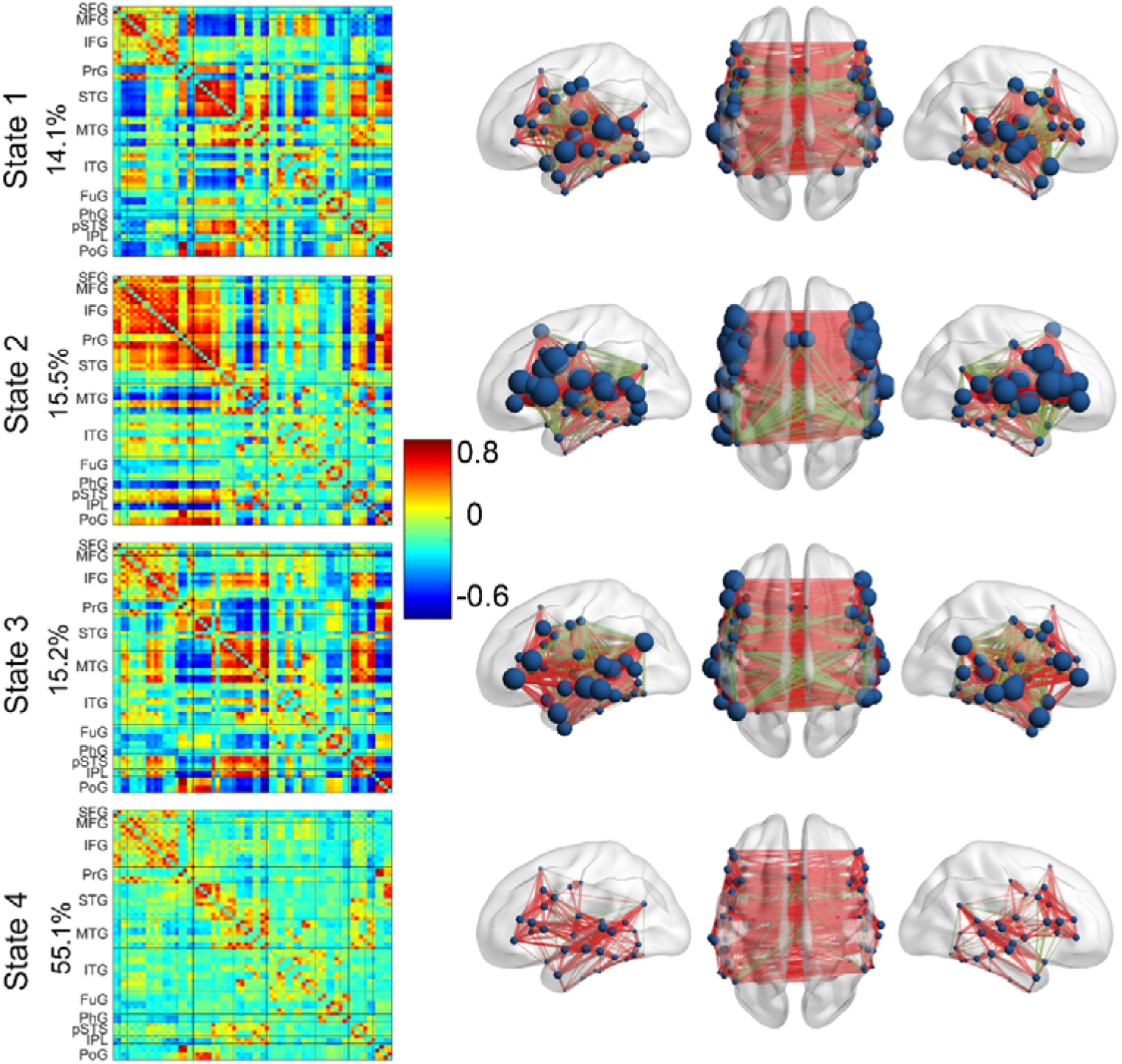
The four temporal reoccurring dFC states and nodal degree distributions. The percentage values represent the overall state frequency. Connections with red color denotes positive connections > 0.2, while connections with green color denotes negative connections < 0.2.

### 3.2 State-dependent hub distributions

We then calculated the nodal degree to assess the hub distribution in each state. Nodes with nodal degree > mean values of the four states were defined as hubs. The hub distributions in the first three states were shown in Figure 3A and Supplementary Table 2. In state 1, 32 nodes were identified as hubs and mainly distributed in STS, pSTS, PoG and PrG. In state 2, 38 nodes were identified as hubs and mainly distributed in prefrontal cortex and posterior temporal gyrus. In state 3, 32 nodes were identified as hubs and mainly distributed in temporal cortex (STG, MTG, pSTS, and IPL) and pars triangular part of IFG. Several hubs in STG and PrG of state 1 were also identified as hubs in state 2. Several hubs in IFG, posterior MTG, and MFG of state 2 were also identified as hubs in state 3. No hubs were found in state 4.

**Figure 3.**
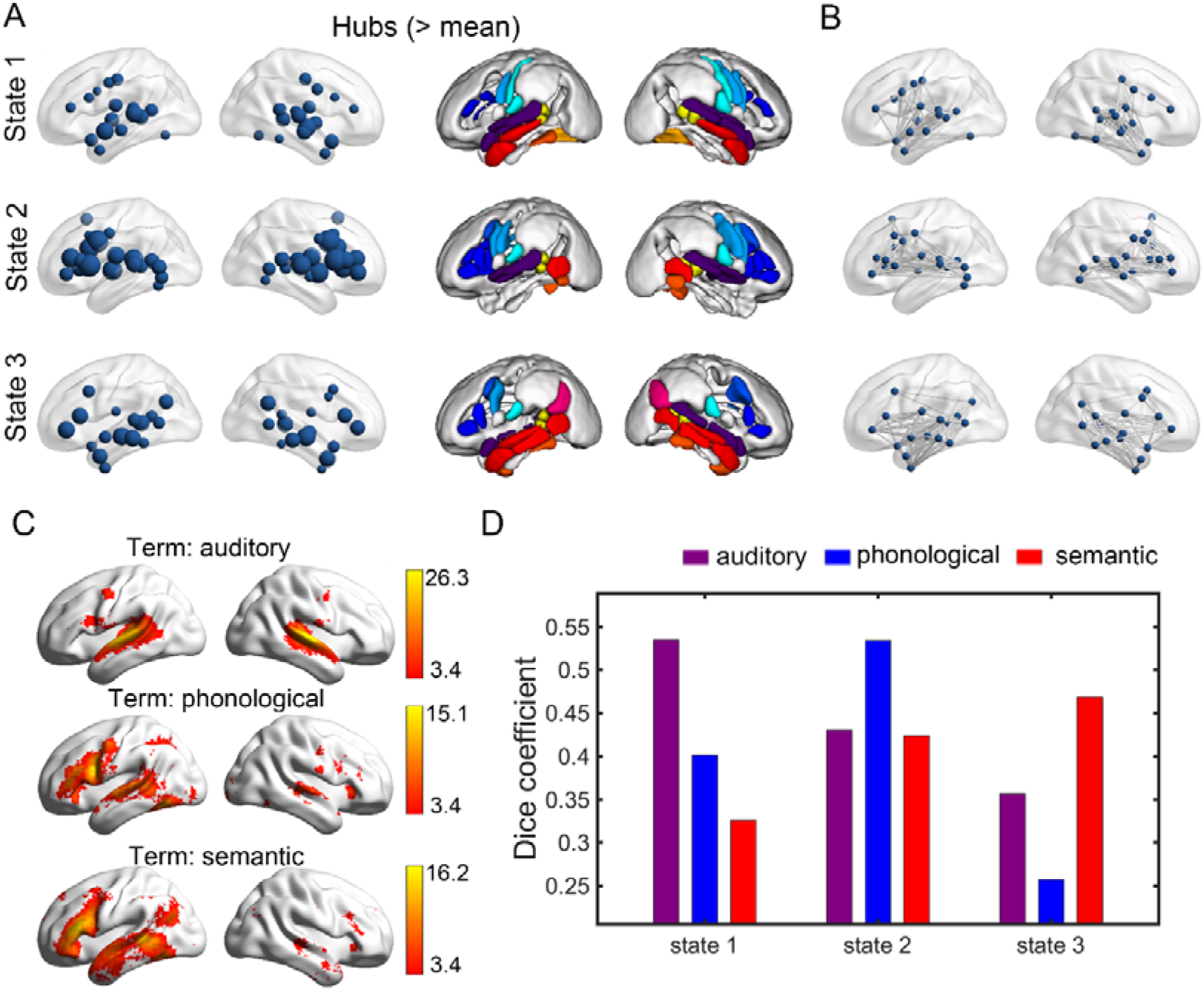
The functional hubs and structural connections of the first three states and the meta-analytic results of auditory, phonological, and semantic processing. A: nodes with strength larger than the mean value of the four states are defined as network hubs. B: the binary white matter connections among hubs were extracted (the whole brain white matter connections are available at https://pan.cstcloud.cn/web/share.html?hash=JX89RKcgTHM). The white matter matrix of the whole language network was shown in the Supplementary Figures 2 A and B. C, the meta results of auditory processing (1252 studies, 46557 activations), phonological processing (377 studies, 17844 activations), and semantic processing (1031 studies, 40030 activations), which are from term-based meta-analyses of association tests with FDR correction (*P* < 0.01, https://www.neurosynth.org/). Neurosynth performs an automated selection of studies based on the predefined term. It divides the entire database of coordinates into two sets: those that occur in articles containing a particular term and those that do not. Then it performs meta-analysis using forward inference maps, which reflect the likelihood that a voxel will activate if a study uses a particular term. D: the dice coefficients between binary images of hub nodes and meta results (dice = 2* (hub * meta)/(hub + meta)). Considering the left-lateralized activations of meta results, the dice coefficients were calculated in the left hemisphere.

The hub distributions showed similar patterns under different correlation thresholds (i.e., *r* > 0.1 and *r* > 0.3) (Supplementary Figure 1).

We also found that hub nodes were densely connected by the underlying white matter fibers (Figure 3B).

To assess the functional relevance of the first three states, we performed term-based meta-analyses at https://www.neurosynth.org/analyses/terms/. We calculated the dice coefficients between the hubs and meta results to assess each state’s functional relevance. According to the hub distributions, three terms, i.e., auditory, phonological, and semantic were selected. State 1 was most likely to be implicated in auditory processing (dice coefficients: auditory, 0.54; phonological, 0.4; semantic, 0.33). State 2 was most likely to be implicated in phonological processing (dice coefficients: auditory, 0.43; phonological, 0.53; semantic, 0.42), and state 3 was most likely to be implicated in semantic processing (dice coefficients: auditory, 0.36; phonological, 0.26; semantic, 0.47) (Figure 3 C and D).

### 3.3 State-dependent modular architectures

To assess the functional segregation of each state, we calculated the modular architecture for each state. The state-dependent modular assignments were shown in Figure 4. In the first three states, the hub distributions highly resembled the corresponding modular assignments. In state 1, two modules were identified. The first module includes almost all nodes in prefrontal cortex and ITG, FuG. The rest nodes were assigned to the second module, which included almost all hubs in this state (26/32=81%). In state 2, three modules were identified. The first module was very similar to the hub distributions of state 2 (26/28=93%). Nodes in the MTG, anterior temporal lobe (ATL), and IPL were in the second module. Nodes in the ITL and FuG were in the third module. Four modules were identified in state 3. The first module includes nodes in prefrontal cortex (SFG, MFG, IFG, and PrG) and posterior ITG. Nodes in the second module include nodes mainly in the temporal cortex (STG, MTG, pSTS, and IPL) and the pars triangular part of IFG, which included almost all hubs in this state (29/32=91). Nodes in STG, PrG, and PoG form the third module. Nodes in ITG, PhG, and FuG form the fourth module. The module assignment in state 4 was a little different from state 3. The pars triangular part of IFG was assigned to the first module (mainly the prefrontal cortex).

**Figure 4.**
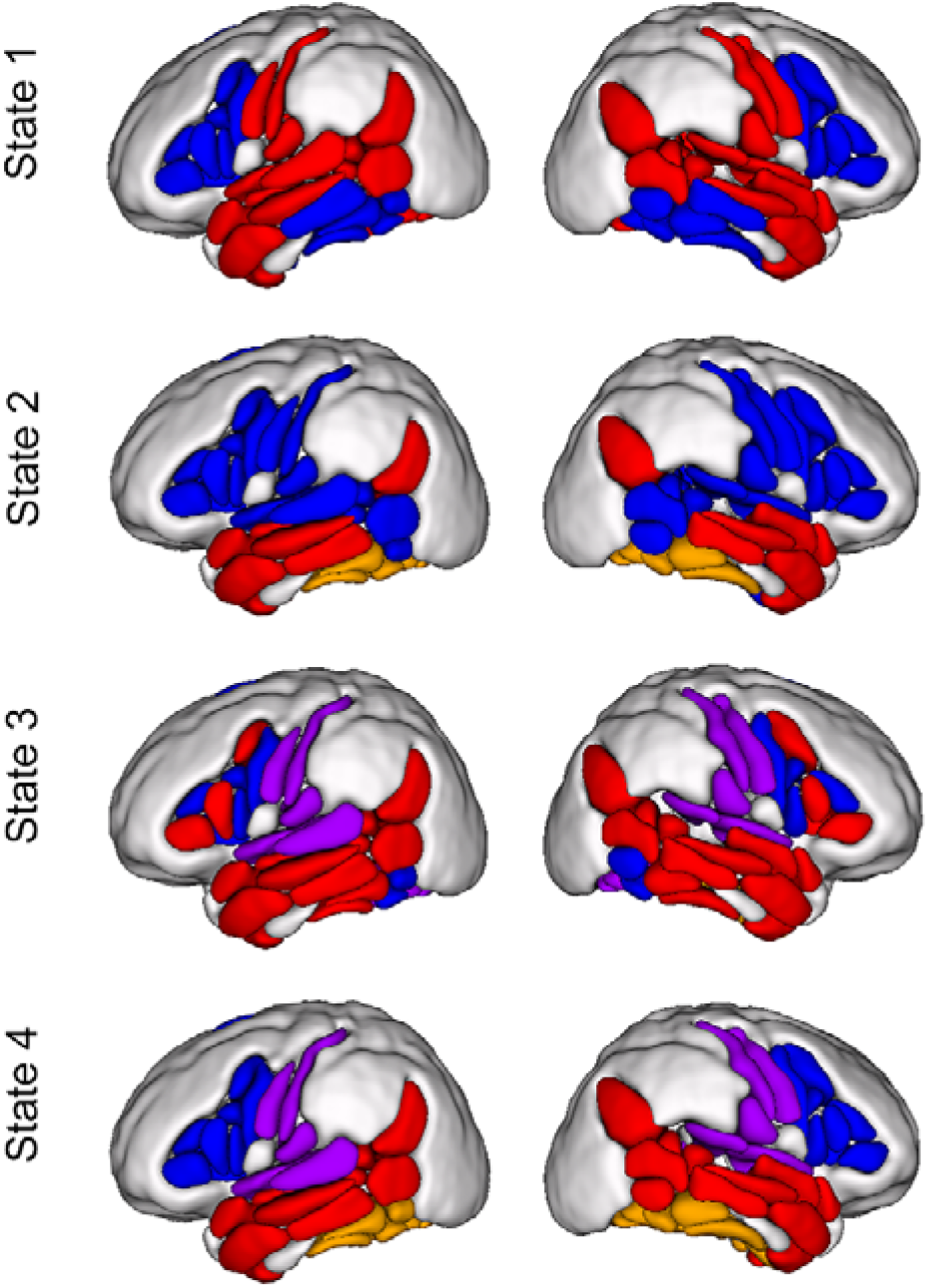
Modular assignments for each state. The module numbers are: in state 1, 2 modules, the red module includes 81% of the hubs; in state 2, 3 modules, the blue module includes 93% hubs; in state 3, 4 modules, the red module includes 91% hubs; in state 4, 4 modules.

### 3.4 Temporal characteristics

Temporally, the first three states appeared exclusively in limited time bins (State 1: 14.1%; state 2: 15.5%; State 3: 15.2%), and for most time (> 55%), the language network kept inactive in state 4. The transition probabilities from state 4 to the first three states were significantly higher than the transition probabilities among the first three states (*Ps* < 0.001, Figure 5).

**Figure 5.**
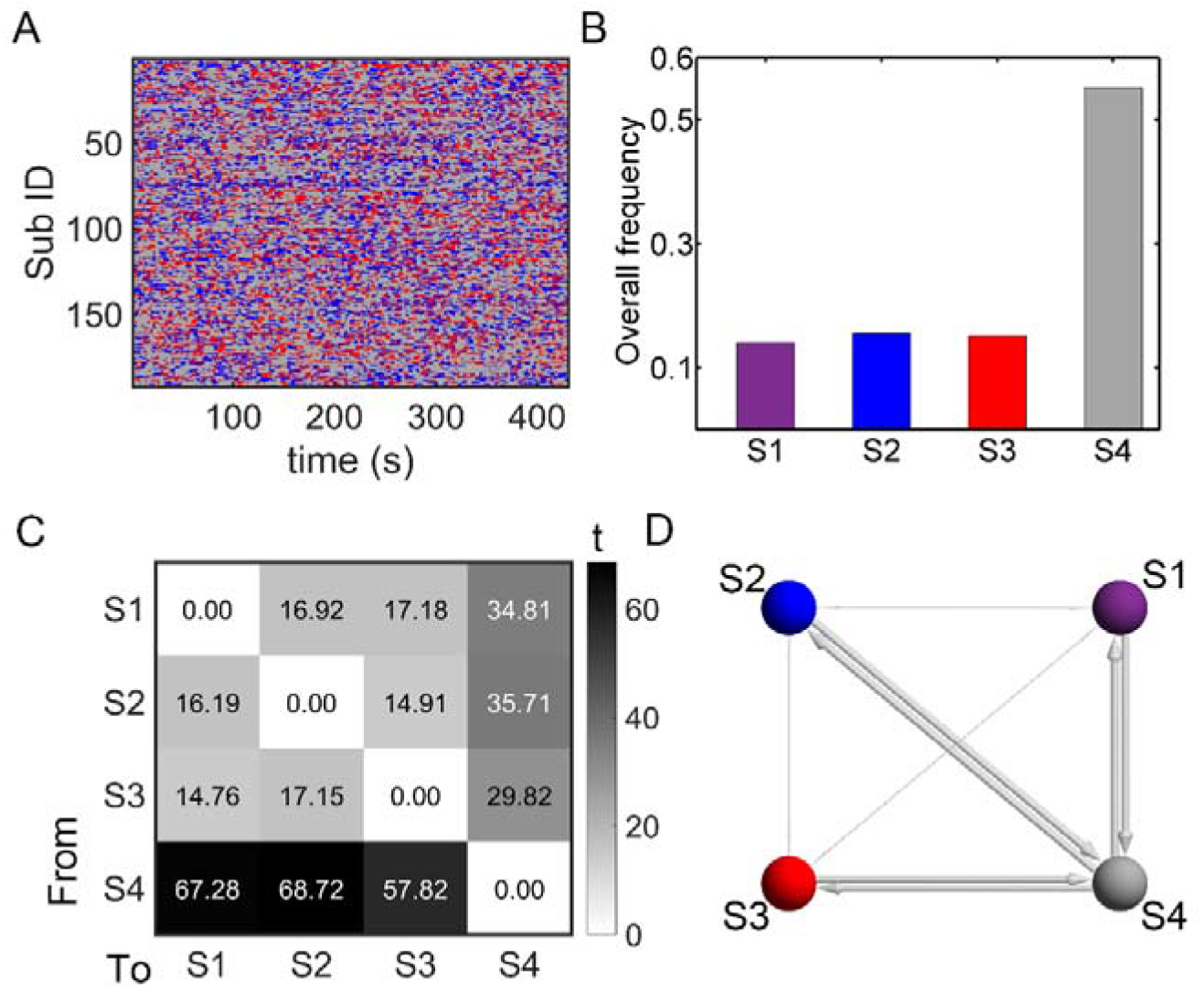
Temporal characteristics of language network dynamics. A: frame-wise state transitions during the scanning (215 volumes, 430 seconds) of each subject (purple, state1; blue, state2; red, state3; gray, state 4). B: overall state frequency, i.e., the percentage of a state of all windows across the group. C: Diagonal-free transition probabilities among states obtained via one-sample t-tests. D: Schematic illustration of the preferred paths (directed). Sphere balls represent states, and line width is proportional to the *t* values.

### 3.5 The spatial characteristics of sFC

The assess similarity and differences of network organization between dFC and sFC, we also analyzed the spatial characteristics of group mean sFC. The connectivity pattern, nodal hubness, and the modular assignment of sFC were shown in the Supplementary Figure 2C and D. Compared to the four dFC states, the sFC showed distinct hub distributions. Nodes within the prefrontal cortex, STG and MTG were highly connected. The modular assignments of sFC were similar to the modular assignments of state 3.

### 3.6 Validation results of whole-brain dynamics

To investigate whether the observed language dynamics were parts of or primarily driven by the whole brain dynamics, we performed whole-brain dFC analyses. In whole-brain level, four temporal reoccurring states were identified (Supplementary Figure 3). The hubness of language areas in whole-brain level was distant from that only calculated in language network.

### 3.7 Validation results of HCP data

In the validation dataset, four temporal reoccurring states were identified. The spatiotemporal characters and state-related hub distributions were highly consistent with the results of the discovery dataset (Supplementary Figures 4 and 5).

### 3.8 DFCs significant predicted individual linguistic performance

To investigate the behavioral relevance of the dFC states, we constructed dFC-linguistics prediction model using the validation dataset. Consistent with the prediction results by Cui et al (2018), the subject median dFCs of state 1 (*r* = 0.24, permutation *P* = 0.01), state 2 (*r* = 0.27, *P* < 0.001), state 4 (*r* = 0.21, *P* = 0.01) and their combinations of the four states (*r* = 0.24, *P* = 0.005) significantly predicted individual ORRT scores (Supplementary Figure 6). The subject median dFCs of state 1 (*r* = 0.23, *P* = 0.006), state 2 (*r* = 0.16, *P* = 0.02), state 3 (*r* = 0.28, *P* = 0.001), state (*r* = 0.24, *P* = 0.001) and the combinations of the four states (*r* = 0.22, *P* < 0.001) significantly predicted individual PVT scores. Meanwhile, sFCs significantly predicted the individual ORRT (*r* = 0.27, permutation *P <* 0.001) and PVT scores (*r* = 0.29, *P* < 0.001).

**Table 1.**
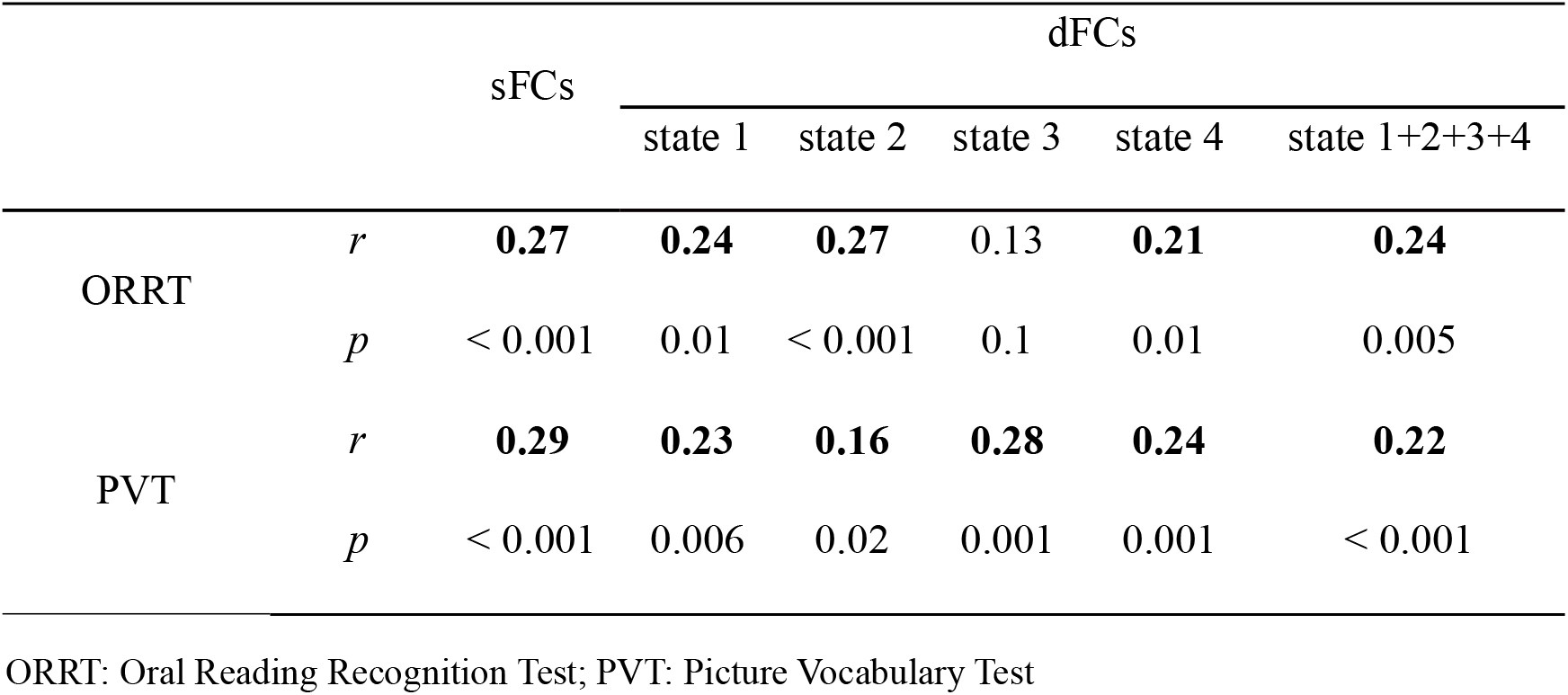
FC-based prediction accuracies for individual language scores of HCP dataset (n = 500)

## 4 Discussion

In this study, we investigated the transient network reorganization underlying the “hierarchy” and “asymmetric” recruitment of the dual-stream model for language and speech processing in the resting-state. We found that the spontaneous brain dynamics of the language network flowed in a domain-separation manner: 1) The framewise language network dynamics were robustly clustered into four low-dimensional states; 2) Hubs in the first three states highly resembled the neurobiology of primary auditory processing and lexical-phonological processing, motor speech production processing, and semantic processing, respectively. While, the fourth state was characterized by the weakest overall functional connectivity but longest state duration (> 55%), which subserved as a baseline state; 3) Behaviorally, machine learning-based dFC-linguistics prediction analyses showed that connections of the four states significantly predicated individual language scores; 4) The dynamic language organization was distinct from the “static” pattern. These results suggested that the hierarchical and parallel architecture of language and speech processing was supported by domain-separation low-dimensional dynamic states, which are essential for flexible functional segregation and integration of various cognitive processes. Based upon these findings, a dynamic “meta-networking” framework of language network in resting state is proposed.

### The first three states and their cognitive function

In state 1, regions in the STG were most active. Except for the phoneme perception in the primary auditory cortex (Bhaya-Grossman and Chang, 2022), the anterior STG is a critical part of auditory ventral stream for word-form recognition, while the middle and posterior STG are critical regions of the auditory dorsal stream for speech monitoring and motor control (DeWitt and Rauschecker, 2013). Moreover, the posterior STG (pSTG) and STS (core areas of Wernicke’s area) also have critical roles in lexical-phonological representation (Binder, 2017). Phoneme perception, word-form recognition, and lexical-phonological representation are primary steps for speech repetition, word and sentence comprehension, and production (Binder, 2017; Hickok, 2022). Term-based meta-analyses further confirmed the functional specificity of state 1 in auditory processing and phonological processing. The independence of state 1 endows the brain to flexibly and dynamically integrated it with the high-level dorsal (phonological) stream (i.e., state 2) or ventral semantic stream (i.e., state 3).

In state 2, hubs are mainly distributed in the prefrontal cortex and posterior temporal cortex. Hub distributions highly resemble the meta results of high-level phonological processing. The pars opercularis of IFG, the ventral part of percental gyrus (vPrG), posterior STG, pSTS, and the pars triangularis of IFG are the most active regions of state 2. The pars opercularis of IFG and vPrG are critical for articulation (Basilakos et al., 2018; Hickok, 2022; Lu et al., 2021). The pars triangularis of IFG has been proposed to process linear sequences of morphemes (Hickok, 2022; Matchin and Hickok, 2020), complex syntactic processes (Friederici and Gierhan, 2013; Wilson et al., 2011), speech planning (Castellucci et al., 2022), and inner speech (Geva et al., 2011). Thus, state 2 may be mainly implicated in motor and speech production processing.

In state 3, hubs mainly include the IPL, MTG, ATL, and IFG, which were well aligned to the neurobiology of semantic processing (Binder et al., 2009; Hodgson et al., 2021; Xu et al., 2016). We also observed that regions involving the anterior STG were also active in state 3. The semantic representation is a core step for word and sentence comprehension, and sentence production (Binder, 2017; Hickok, 2022). Moreover, semantic processing is also central to thought (Buckner and DiNicola, 2019), object recognition, and use (Patterson et al., 2007). Besides the critical role in language and speech processing, it is also tightly related to memory and sensorimotor systems. The independence of state 3 endows the brain to flexibly and dynamically retrieved or manipulated concept knowledge during various cognitive processes.

Several regions (e.g., IFG, pSTG, pSTS, and pMTG) were active in more than one state, which was in line with their functionally heterogeneous roles and the overlap representations of phonological processing, semantic processing, and syntactic processing in these regions (Blumstein and Amso, 2013; Chen et al., 2021; Friederici and Gierhan, 2013; Hodgson et al., 2021; Wang et al., 2020).

The time-varying, dynamic changes in functional connectivity in tasks are time-locked to task state and performance (Cohen, 2018; Shine and Poldrack, 2018). Nevertheless, it is hard to speculate the roles of these endogenous regenerated domain-specific states in verbal mind-wandering. In the resting-state, several spontaneous thoughts, such as imagined and intended speech (flexibly manipulating verbal structures to express dynamically changing mental representations) (Proix et al., 2022) or flexibly construct representations of others’ intended meanings (Deak, 2003) are language-related (Alderson-Day and Fernyhough, 2015; Lœvenbruck et al., 2018). It has been suggested that inner speaking and inner hearing activities are conceived of as multimodal acts with multisensory percepts, and also need dynamically integrate diverse cognitive networks (e.g., default mode network, somatosensory network) into a coherent pattern (Doucet et al., 2012). Hence one alternative explanation of the spontaneous dynamics of language network is that they may be preconfigured in a domain-separation manner that affords rapidly integrated with other networks to flexibly and continuously explore an array of available cognitive architectures for inner or external language and speech tasks (Gonzalez-Castillo and Bandettini, 2018; Pezzulo et al., 2021).

### The structural bases of language network dynamics

We also showed that the spontaneous brain dynamics of language network were directly constrained by the underlying white matter connections. White matter connections would directly shape the strength, persistence, and spatial statistics of spontaneous brain dynamics (Cabral et al., 2017; Fukushima et al., 2018; Honey et al., 2009). The middle longitudinal fascicle is a recently delineated association cortico-cortical fiber pathway in humans. It courses through STG and IPL connecting STG and temporal pole principally with the angular gyrus, which is consistent with the hub distributions in state 1 (de Champfleur et al., 2013; Makris et al., 2009; Maldonado et al., 2013; Wang et al., 2013). The arcuate fasciculus (AF) is the major white matter of the dorsal stream of language model, which connects the posterior parts of IFG to the middle and posterior parts of STG and MTG, and participates in phonological and motor processing for speech output (Dick et al., 2014; Glasser and Rilling, 2008; Saur et al., 2008; Yagmurlu et al., 2016; Zhang et al., 2021). Moreover, the AF projects beyond the superior temporal gyrus in the temporal lobe and also includes middle, inferior, and temporobasal fibers (Giampiccolo and Duffau, 2022), which are consistent with the hub distributions in state 2. The ventral (semantic) stream, which involving multi association tracts, including the inferior fronto-occipital fasciculus (connecting occipital cortex, superior parietal lobule, ITG, and IFG), inferior longitudinal fasciculus (connecting occipital and temporal cortices) and uncinate fasciculus (connecting the ATL and IFG) are consistent with the hub distributions in state 3.

### A dynamic “meta-networking” framework of the language network in the resting-state

We synthesized all the spatiotemporal properties and proposed a dynamic “meta-networking” framework for the language network in the resting-state (Figure 6). The dynamic framework is in line with the dual-stream model of speech and language processing. It includes four low-dimensional states, which flow in a domain-separation manner. According to the hub distributions, state 1 is mainly implicated in phoneme perception and lexical-phonological processing, state 2 is mainly implicated in motor and speech production processing, and state 3 is the semantic system. These “meta states” are directly constrained by the underlying white matter connectivity, and can also significantly predict individual language performance during tasks. The dynamic “meta-networking” framework can subserve the network bases for flexible and hierarchical functional segregation and integration during language and speech processing.

**Figure 6.**
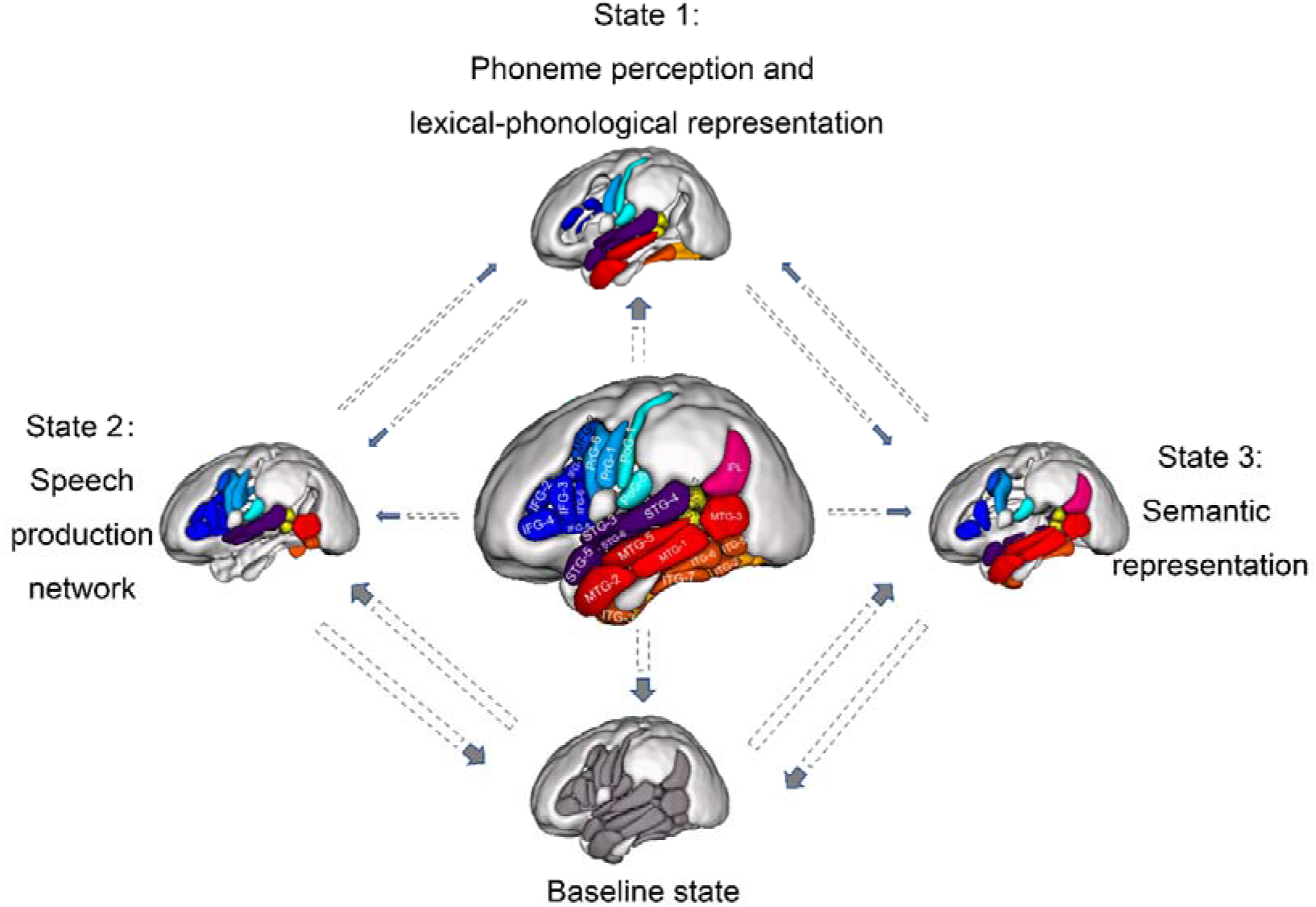
The dynamic “meta-networking” framework of language network in the resting-state. The domain-separation language network dynamics are spatially constrained by the underlying white matter connectivity, have distinct temporal and spatial characters, are in line with the dual-stream model of speech and language processing, have cognitive and behavioral relevance, and subserve the network bases for flexible and hierarchical functional segregation and integration during language and speech processing. The sizes of the arrays are proportional to the statistical state transition probabilities. Note that several nodes are active (i.e., hub) in multi states, e.g., pMTG, pars triangularis of IFG, and pSTS in state 2 and 3. These findings are in line with their functionally heterogeneous roles and the overlap representations of phonological and semantic processing in these regions.

### The clinical relevance of the dynamic “meta-networking” framework

Different from those dynamic studies focusing on the whole-brain network, our results revealed the fine-grained low-dimensional language network dynamics in the resting-state. These states may have clinical relevance to better understanding the neuroimaging alterations underlying different types of language or speech deficits. Firstly, these cognition-related states have the potential to be effective biomarkers for language or speech deficits associated with expressive and receptive language or speech skills (American Psychiatric Association and Association, 2013). Second, the dynamic framework of language network could subserve as a “meta-networking” (network of networks) (Duffau, 2021), which would be useful for mapping and distinguishing subtypes of patients with different language or speech deficits severity (Bonkhoff et al., 2020; Guo et al., 2019; He et al., 2018), degree of neuroplasticity (Liu et al., 2020; Yuan et al., 2020) and recovery trajectories (Yuan et al., 2022).

## 5 Conclusions

Frame-wise dynamic functional connectivity analyses hold promise to refine the temporal network organization (meta-networking) of functional-specific networks. We found that the time-varying functional connectivity of language network in the resting-state is temporally and spatially non-random but robustly clustered into four low-dimensional states. These states are spatially constrained by the underlying white matter connectivity, have distinct temporal and spatial characters, are in line with the dual-stream model of language network, and have cognitive relevance. These findings hint a domain-separation manner of language network dynamics in the resting-state, forming a dynamic “meta-networking” framework, which may be effective biomarkers for diagnosing different subtypes of aphasia and detecting the language or speech recovery.

## Data/code availability statement

The discovery dataset was part of the 1000 Functional Connectomes Project and is available at http://fcon_1000.projects.nitrc.org. The validation dataset was from the Human Connectome Project (HCP) 900 Subject Release (see Van Essen et al., 2012 for dataset details), which is available at http://www.humanconnectome.org/. The DCC toolbox was available at https://github.com/canlab/Lindquist_Dynamic_Correlation/tree/master/DCC_toolbox. Other software used in the study was based on DPABI (http://rfmri.org/dpabi), SPM (http://www.fl.ion.ucl.ac.uk/spm), GRETNA (https://www.nitrc.org/projects/gretna/) and BrainNetViewer (https://www.nitrc.org/projects/bnv/).

## CRediT authorship contribution statement

**Binke Yuan**: Conceptualization, Methodology, Formal analysis, Investigation, Writing – review & editing, Writing – original draft, Visualization. **Hui Xie**: Conceptualization, Methodology, Formal analysis, Investigation, Writing – review & editing, Writing – original draft, Visualization. **Zhihao Wang**: Conceptualization, Methodology, Formal analysis, Investigation, Writing – review & editing, Writing – original draft, Visualization. **Yangwen, Xu**: Conceptualization, Investigation, Writing – original draft. **Hanqing Zhang**: Conceptualization, Methodology, Formal analysis. **Jiaxuan Liu**: Conceptualization, Methodology, Formal analysis. **Lifeng Chen**: Conceptualization, Methodology, Formal analysis. **Chaoqun, Li**: Conceptualization, Methodology, Formal analysis. **Shiyao Tan**: Conceptualization, Methodology, Formal analysis. **Zonghui Lin**: Conceptualization, Methodology, Formal analysis. **Xin Hu**: Conceptualization, Methodology, Formal analysis. **Tianyi Gu**: Conceptualization, Methodology, Formal analysis. **Junfeng Lu**: Conceptualization, Methodology, Formal analysis. **Dongqiang Liu**: Conceptualization, Methodology, Formal analysis, Investigation, Writing – review & editing, Writing – original draft, Visualization. **Jinsong Wu**: Conceptualization, Methodology, Formal analysis, Investigation, Writing – review & editing, Writing – original draft, Visualization.

## Funding

The study is supported by the National Social Science Foundation of China (No. 20&ZD296), Key-Area Research and Development Program of Guangdong Province (No. 2019B030335001), National Natural Science Foundation of China (No.32100889). The funding agencies took no part in the design or implementation of the research.

## Conflict of Interest Statement

The authors declare no competing financial interests.

## Notes

### Competing Interest Statement

The authors have declared no competing interest.

