## SUPPLEMENTARY MATERIALS for "The domain-separation low-dimensional language network dynamics in the resting-state support the flexible functional segregation and integration during language and speech processing"

**Supplementary Methods**

**1 Preprocessing steps of HCP data**

It comprised the following steps: 1) discarding the first 50 volumes to decrease the signal’s instability; 2) realignment; 3) co-registering the T1-weighted image to the corresponding mean functional image; 4) segmenting into gray matter, white matter, and cerebrospinal fluid by DARTEL; 5) regressing common nuisance out by Compcor (Behzadi et al., 2007), including the white matter signal, the cerebrospinal fluid signal, 24 movement regressors, and global signal. The 24 movement regressors included autoregressive models of motion incorporating 6 head motion parameters, 6 head motion parameters one time point before, and the 12 corresponding squared items (Yan et al., 2013); 6) detrending; 7) normalizing to the standard Montreal Neurological Institute space (MNI template, resampling voxel size 3 × 3 × 3 mm3); 8) smoothing with a Gaussian kernel of 4 mm full width at half maximum; and 9) filtering (0.01–0.1 Hz).

**2 The task descriptions, scoring process, and interpretations of ORRT and PVT**

Note that a full version of NIH Toolbox (Scoring and Interpretation Guide) is available in <https://www.epicrehab.com/epic/documents/crc/crc-201307-nih-toolbox-scoring-and-interpretation-manual%209-27-12.pdf>

**2.1 NIH Toolbox Oral Reading Recognition Test (ORRT)**

**Description**: Separate but parallel reading tests have been developed in English and Spanish. In either language, the participant is asked to read and pronounce letters and words as accurately as possible. The test administrator scores them as right or wrong. For the youngest children, the initial items require them to identify letters (as opposed to symbols) and to identify a specific letter in an array of four symbols. The test is given via CAT and requires approximately three minutes. This test is recommended for ages 7–85, but is available for use as young as age 3, if desired.

**Scoring Process**: IRT (Item Response Theory) is used to score the Reading Test. A theta score is calculated for each participant, representing the relative overall reading ability or performance of the participant. Age-Adjusted, Fully Adjusted and Unadjusted Scale Scores as well as a national percentile rank corresponding to the ageadjusted scale score are provided for the Reading Test. In addition, the theta score is converted to a Computed Score, which ranges from roughly 500 to 2500 and can be used for simple reading ability

comparisons over time.

**Interpretation**: The Reading Test is a measure of reading decoding skill and of crystallized abilities, those abilities that are generally more dependent upon past learning experiences and consistent across the life span. To interpret individual performance, one can evaluate all three types of scale scores plus the national percentile; higher scores indicate better reading ability. The Reading Computed Score can be useful in evaluating pure change in performance from one assessment to another. For example, a higher Computed Score for Reading would mean that the participant is able to correctly identify more difficult words on the subsequent assessment, which may indicate developmental growth or a return to a previous level of functioning. Such an interpretation is not norm-referenced, of course. Each language version of the Reading test is calibrated independently using language-specific items administered to a language-specific cohort. Therefore, Reading computed scores are not compatible between different languages (similar scores on the English and Spanish reading tests are not comparable). Note further that reading decoding skill is generally considered easier in the Spanish language than it is in English due to the presence of fewer linguistic exceptions.

**2.2 NIH Toolbox Picture Vocabulary Test (PVT)**

**Description**: This measure of receptive vocabulary is administered in a computerized adaptive format. That is, the next question a participant receives depends on his/her response to the previous question; Computer Adaptive Testing (CAT) ensures a test that is tailored to the participant’s needs. The respondent is presented with an audio recording of a word and four photographic images on the computer screen and is asked to select the picture that most closely matches the meaning of the word. This test takes approximately four minutes to administer and is recommended for ages 3-85.

**Scoring Process**: IRT is used to score the TPVT. A score known as a theta score is calculated for each participant; it represents the relative overall ability or performance of the participant. A theta score is very similar to a z-score, which is a statistic with a mean of zero and a standard deviation of one. Age-Adjusted, Fully Adjusted and Unadjusted Scale Scores, as well as a national percentile rank that corresponds to the age-adjusted scale score, are provided for the TPVT. In addition, the theta score is converted to a “Computed Score” that appears in the Assessment Scores output file available through Assessment Center. (This file and its components are described in detail in Appendix B1.) The Computed Score for TPVT ranges from roughly 200 to 2000 and can be used for simple vocabulary ability comparisons over time.

**Interpretation**: The PVT is a measure of general vocabulary knowledge and is considered to be a strong measure of crystallized abilities (those abilities that are more dependent upon past learning experiences and are consistent across the life span). To interpret individual performance, one can evaluate all three types of scale scores. A participant’s age-adjusted scale score at or near 100 indicates vocabulary ability that is average for the age level. Scores around 115 suggest above-average vocabulary ability, while scores around 130 suggest superior ability – in the top 2 percent nationally for age, based on Toolbox normative data. Conversely, a score of 85 suggests below-average vocabulary ability, while a score in the range of 70 or below suggests significant impairment in language ability, which may also be indicative of difficulties in school (for children) or trouble functioning in work environments with a language demand. An unadjusted scale score allows us to view the participant’s performance in comparison to the entire Toolbox national sample, allowing for a more absolute view of the participant’s ability. The fully adjusted scale scores have been statistically adjusted to level the playing field interpretively, such that an individual’s score can be compared to a narrower group, more similar demographically. The PVT Computed score provides a way of gauging raw improvement or decline from Time 1 to Time 2 (or subsequent assessments). Such a score is useful because a raw score does not provide relevant information on a computer-adaptive test. (Raw scores are useful for monitoring absolute improvement/decline over time when statistical transformations are not used in the scoring process, 6 such as occur in IRT-based scoring or in the Flanker or DCCS measures, described below.) Thus, a computed score of 600 at Time 1 and 640 at Time 2 represents real improvement by the participant in vocabulary knowledge; however, this individual’s Age-Adjusted Scale Score may or may not have increased, depending on how his/her performance at Time 1 and Time 2 compared to the age cohorts used in the national norms. An individual who has made small gains in overall knowledge may still have regressed when compared with age-similar peers if the national sample of peers made larger gains in knowledge over the same period. Thus, one can see the value of the variety of Toolbox scores provided. Each language version of the TPVT is calibrated independently using language-specific items administered to a language specific cohort. Therefore, PVT computed scores are not compatible between different languages (similar scores on the English and Spanish picture vocabulary tests are not comparable).

**Supplementary Figures**

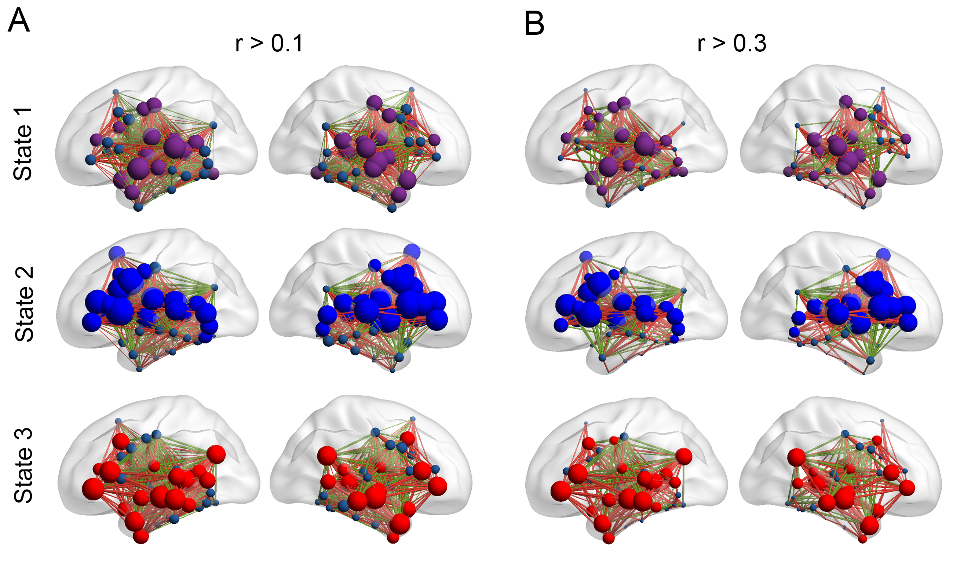

**Supplementary Figure 1.** State hub distributions under different correlation thresholds (i.e., *r* > 0.1 and *r* > 0.3).

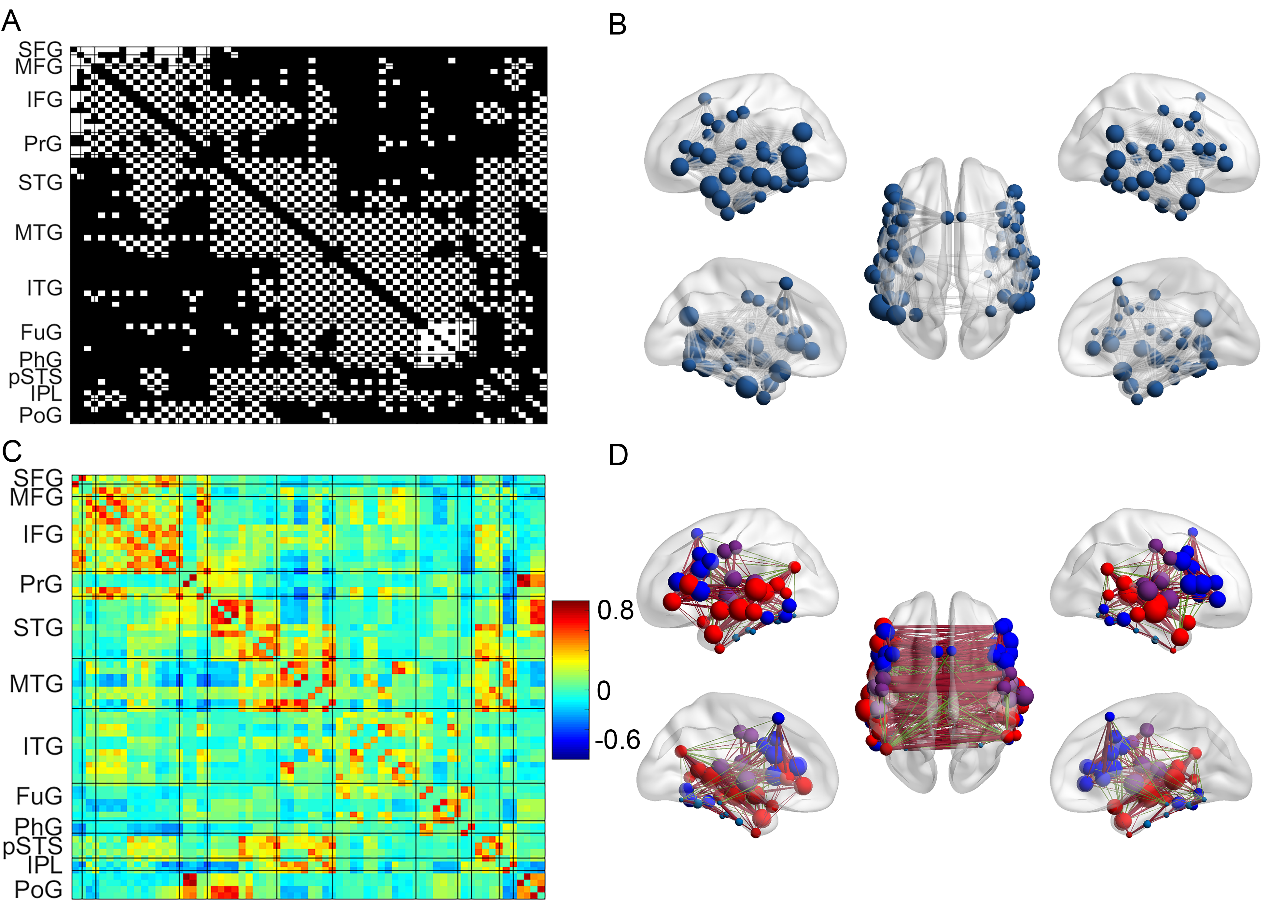

**Supplementary Figure 2.** The matrices of structural and static functional networks for language and speech processing, their corresponding nodal degree distributions, and the modular assignment of the functional network. White squares (A) or gray lines (B) denote the white matter connection between two nodes. In D, connections with dark red color denotes positive connections > 0.2, while connections with blue color denotes negative connections < -0.2. Almost all nodes were densely connected in both the structural and static functional language networks. Four modules were identified in sFC, which were different from the modular assignments of the four states. The first module includes nodes in prefrontal cortex (SFG, MFG, IFG and PrG) and posterior ITG, while the second module includes nodes mainly in temporal cortex (STG, MTG, pSTS and IPL) and triangular part of IFG. Nodes in STG, PrG and PoG form the third module. Nodes in ITG, PhG and FuG form the fourth module.

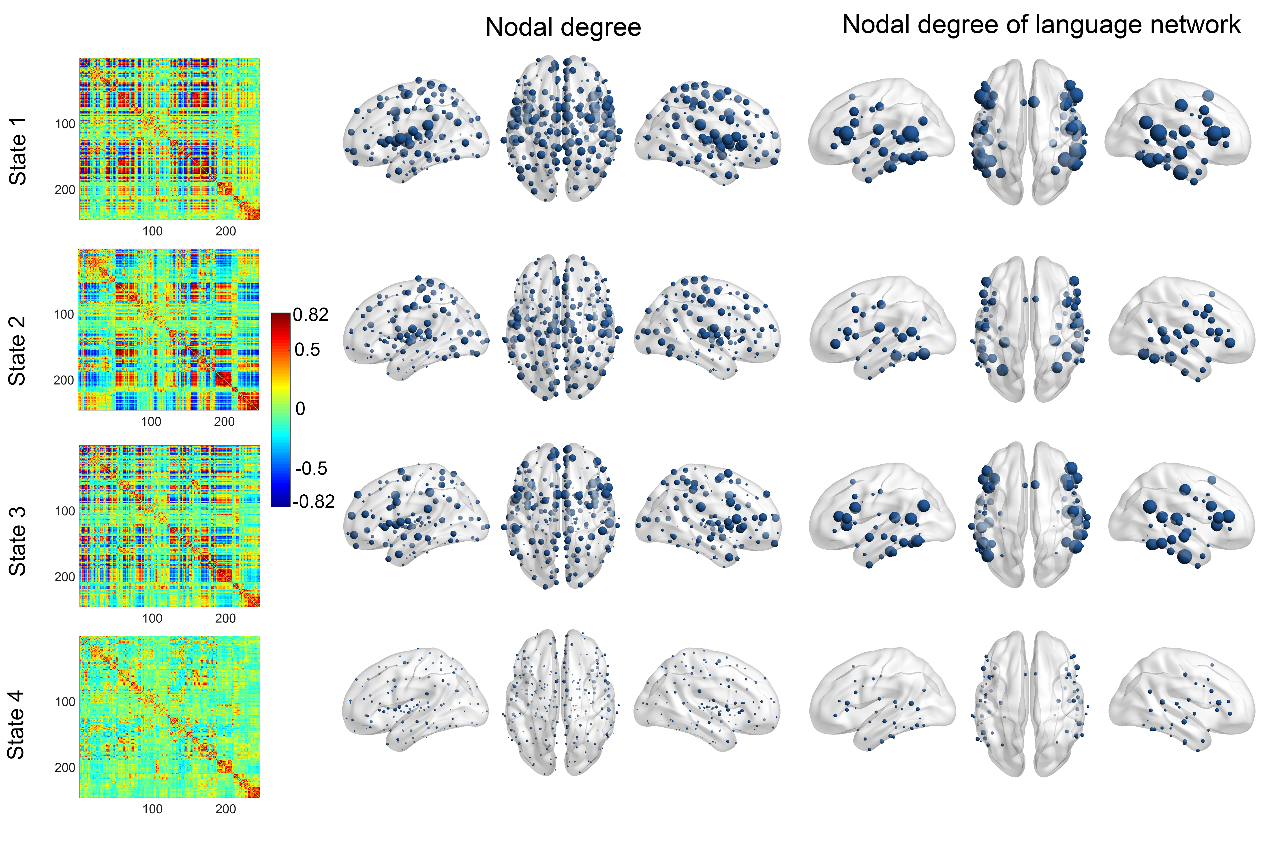

**Supplementary Figure 3.** The whole brain dynamics of Human Brainnetome atlas (n = 246) and state-specific nodal degrees. Nodes belong to the language-speech network were showed again in the right panel.

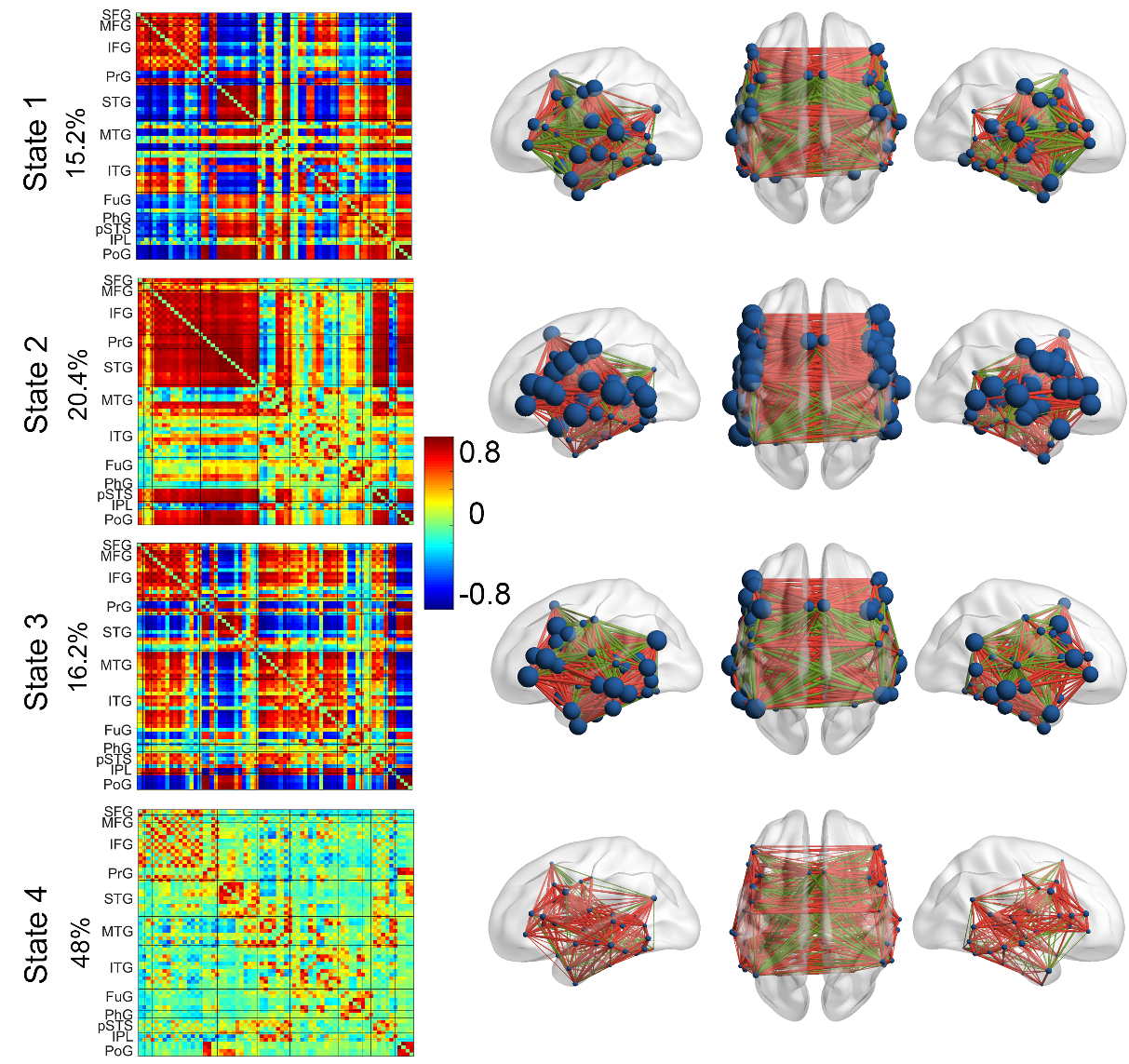

**Supplementary Figure 4.** Validation results of HCP data. The four temporal reoccurring dFC states and nodal degree distributions were shown. The percentages values represent the state's overall frequency. Connections with red color denotes positive connections > 0.2, while connections with green color denotes negative connections < 0.2.

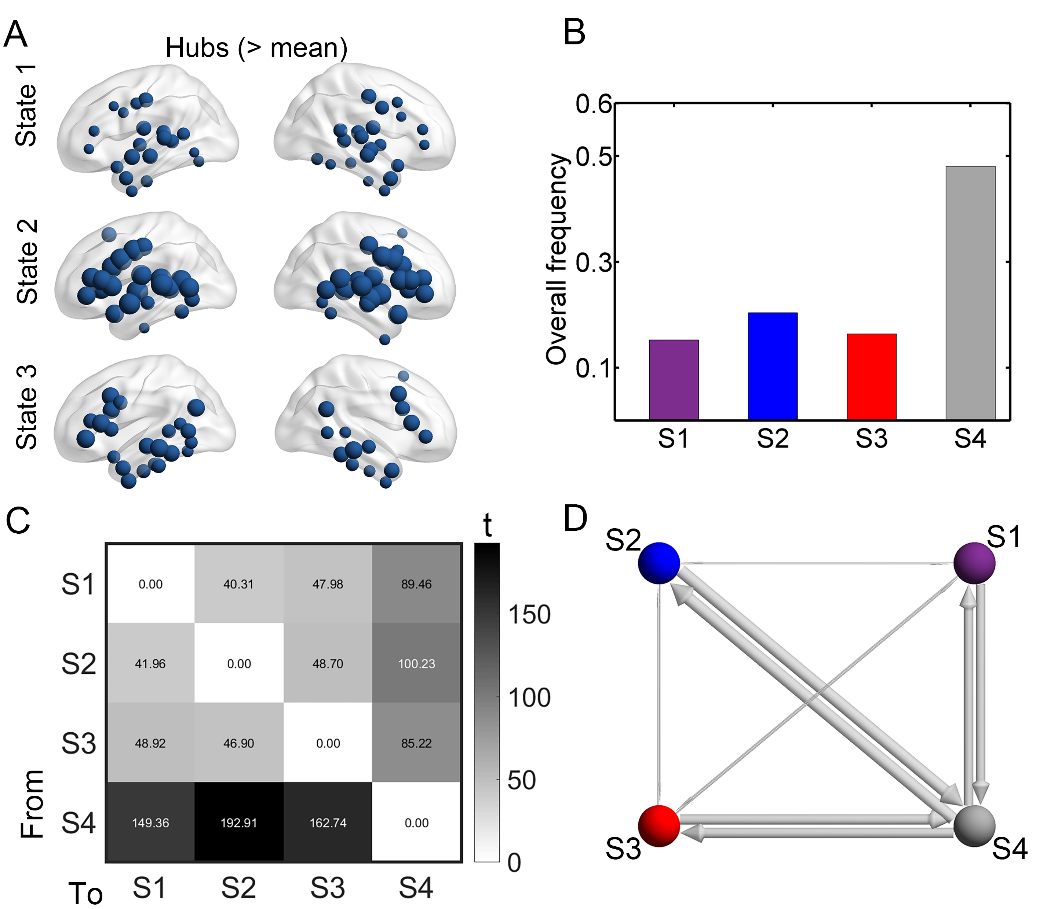

**Supplementary Figure 5.** Spatiotemporal characteristics of HCP data. A: state-related hub (> mean) distributions. No hub was found in state 4. B: state overall frequency, i.e., the percentage of a state of all windows across the group. C: Diagonal-free transition probabilities among states obtained via one-sample t-tests. D: Schematic illustration of the preferred paths (directed). Sphere balls represent states, and line width is proportional to the *t* values.

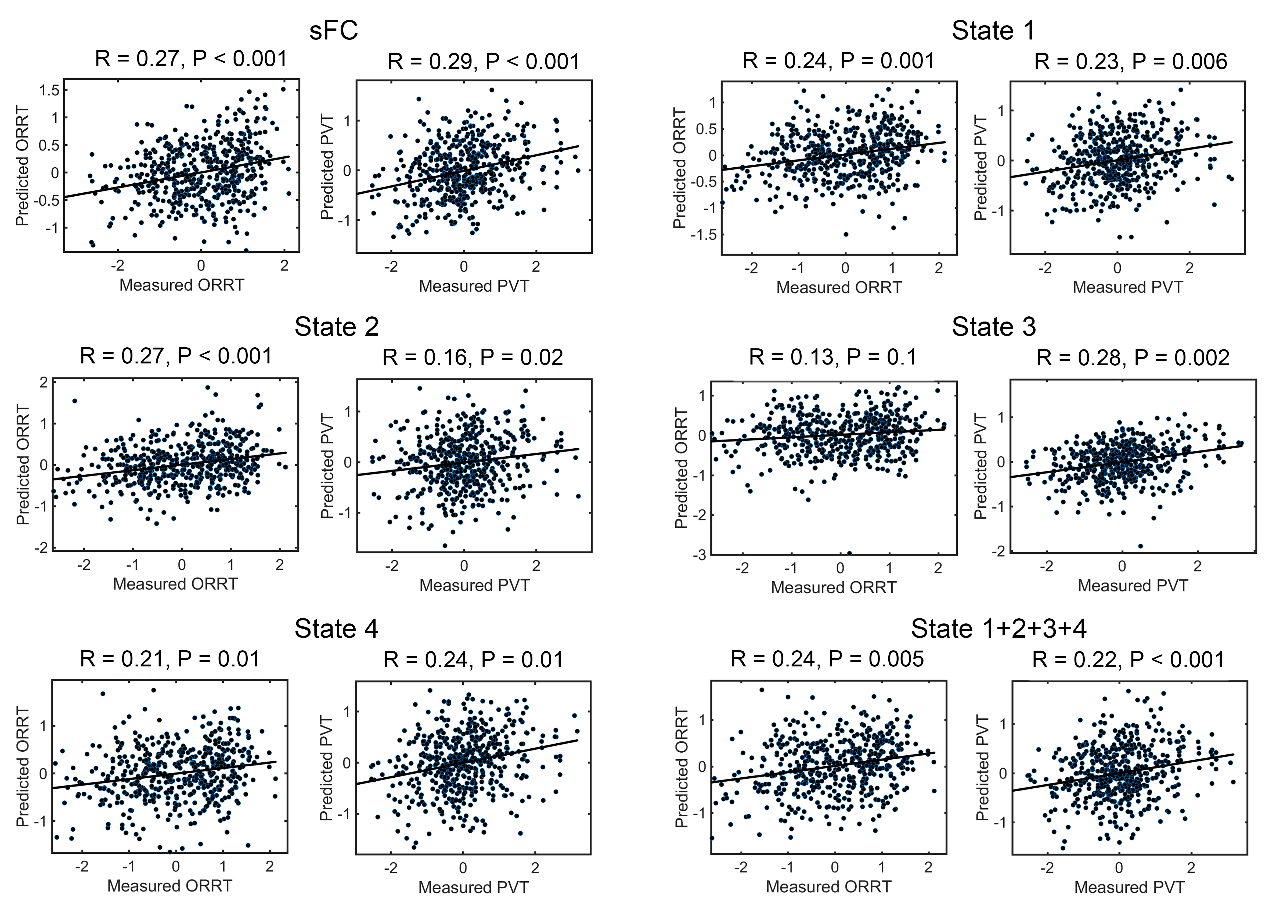

**Supplementary Figure 6.** The FC-language model accuracies and significances (*n* = 500). The scatter plots showed real (normalized within each group) and predicted scores and the corresponding linear fitted lines. The *R* value was the Pearson correlation coefficients between the predicted values and the actual values, the model significance and the *P* values were based on 1000 permutation tests.

**Supplementary tables**

Supplementary Table 1 Anatomical regions，language-related behavioral domains, and paradigm classes of the language network. Coordinates are in the standard Montreal Neurologic Institute Space.

| ID | x | y | z | anatomy | behavioral domains | paradigm classes |
| --- | --- | --- | --- | --- | --- | --- |
| 1 | -4.6 | 15.7 | 53.2 | SFG | Execution. Speech, Phonology, Semantics, Speech | Word Generation (Covert and Overt) |
| 2 | 7.0 | 16.4 | 54.7 | SFG | Homolog of node 1 | - |
| 17 | -41.9 | 13.6 | 36.4 | MFG | Phonology, Semantics | Semantic. Monitor/Discrimination, Word Generation (Covert) |
| 18 | 42.2 | 11.9 | 38.4 | MFG | Homolog of node 17 | - |
| 29 | -45.6 | 13.0 | 23.4 | IFG | Phonology, Semantics, Speech, and Syntax | Phonological. Discrimination, Semantic. Monitor/Discrimination |
| 30 | 45.0 | 15.7 | 25.0 | IFG | Homolog of node 29 | - |
| 31 | -47.6 | 31.6 | 13.6 | IFG | Phonology, Semantics, Speech, and Syntax | Phonological. Discrimination, Semantic. Monitor/Discrimination, Word Generation (Covert and Overt) |
| 32 | 47.8 | 35.1 | 13.2 | IFG | Homolog of node 31 | - |
| 33 | -52.2 | 22.5 | 11.4 | IFG | Semantics, Speech, and Syntax | Reading (Covert), Semantic. Monitor/Discrimination, Word Generation (Covert and Overt) |
| 34 | 54.2 | 23.9 | 11.8 | IFG | Homolog of node 33 | - |
| 35 | -49.1 | 36.3 | -2.8 | IFG | Semantics, Speech, and Syntax | Semantic. Monitor/Discrimination, Word Generation (Covert) |
| 36 | 50.8 | 36.8 | -0.7 | IFG | Homolog of node 35 | - |
| 37 | -39.5 | 22.9 | 3.7 | IFG | Phonology, Semantics, Speech, and Syntax | Semantic. Monitor/Discrimination, Word Generation (Covert) |
| 38 | 42.1 | 22.0 | 3.2 | IFG | Homolog of node 37 | - |
| 39 | -51.3 | 13.2 | 6.0 | IFG | Phonology, Semantics, Speech | Music. Comprehension/Production, Recitation/Repetition. (Covert), Word Generation (Covert) |
| 40 | 53.6 | 14.3 | 11.8 | IFG | Homolog of node 39 | - |
| 53 | -49.5 | -7.1 | 38.8 | PrG | Execution. Speech | Reading (Overt), Recitation/Repetition. (Overt) |
| 54 | 54.9 | -2.0 | 33.3 | PrG | Execution. Speech | Reading (Overt), Recitation/Repetition. (Overt) |
| 63 | -49.1 | 4.7 | 30.5 | PrG | Orthography, Phonology, Semantics, Speech and Syntax | Phonological. Discrimination, Reading (Covert) |
| 64 | 51.1 | 7.2 | 30.9 | PrG | Homolog of node 63 | - |
| 71 | -53.7 | -32.1 | 12.4 | STG | Execution. Speech, Phonology, and Speech | Music. Comprehension/Production, Passive Listening, Phonological. Discrimination, Reading (Overt), Recitation/Repetition. (Covert and Overt) |
| 72 | 54.5 | -23.7 | 10.6 | STG | Execution. Speech, Phonology | Music. Comprehension/Production, Passive Listening, Phonological. Discrimination, Recitation/Repetition. (Overt) |
| 73 | -50.1 | -10.3 | 1.1 | STG | Execution. Speech, Phonology, Speech | Music. Comprehension/Production, Passive Listening, Phonological. Discrimination, Reading (Overt), Recitation/Repetition. (Overt) |
| 74 | 51.1 | -3.7 | -0.9 | STG | Execution. Speech, | Music. Comprehension/Production, Passive Listening, Recitation/Repetition. (Overt) |
| 75 | -62.8 | -32.8 | 7.4 | STG | Execution. Speech, Phonology, Semantics, Speech | Passive Listening, Phonological. Discrimination, Reading (Overt), Semantic. Monitor/Discrimination |
| 76 | 66.5 | -20.8 | 6.6 | STG | Execution. Speech, Phonology, Speech | Music. Comprehension/Production, Passive Listening, Phonological. Discrimination, Reading (Overt) |
| 77 | -45.1 | 10.5 | -19.4 | STG | Homolog of node 78 | - |
| 78 | 47.1 | 12.3 | -19.7 | STG | Speech | Film Viewing, Passive Listening |
| 79 | -55.1 | -3.2 | -10.1 | STG | Execution. Speech, Phonology, Semantics, Speech | Music. Comprehension/Production, Passive Listening, Phonological. Discrimination, Reading (Overt) |
| 80 | 55.8 | -12.5 | -5.2 | STG | Execution. Speech, Phonology, Semantics, Speech | Music. Comprehension/Production, Passive Listening, Phonological. Discrimination, Reading (Overt), Semantic. Monitor/Discrimination |
| 81 | -65.2 | -30.9 | -11.3 | MTG | Semantics | Semantic. Monitor/Discrimination |
| 82 | 64.5 | -29.2 | -13.2 | MTG | Homolog of node 81 | - |
| 83 | -53.2 | 2.2 | -29.6 | MTG | Language | Semantic. Monitor/Discrimination, Reading (Covert) |
| 84 | 51.1 | 5.7 | -31.8 | MTG | Language | Passive Listening |
| 85 | -58.9 | -57.6 | 4.3 | MTG | Semantics and Syntax | Semantic. Monitor/Discrimination, Word Generation (Overt) |
| 86 | 60.1 | -53.3 | 2.9 | MTG | Homologues of node 85 | Film Viewing |
| 87 | -58.5 | -19.8 | -9.4 | MTG | Phonology, Semantics, Speech, and Syntax | Passive Listening, Phonological. Discrimination, Reading (Covert), Semantic. Monitor/Discrimination |
| 88 | 58.3 | -15.4 | -10.1 | MTG | Semantics and Speech | Passive Listening, Phonological. Discrimination |
| 89 | -45.5 | -26.7 | -26.1 | ITG | Orthography and Semantics | Reading (Covert), Semantic. Monitor/Discrimination |
| 90 | 45.8 | -14.6 | -32.4 | ITG | Homolog of node 89 | - |
| 91 | -50.5 | -57.0 | -14.1 | ITG | Phonology and Semantics | Naming (Overt) |
| 92 | 53.5 | -52.4 | -18.5 | ITG | Phonology and Semantics | - |
| 93 | -43.7 | -2.9 | -41.4 | ITG | Semantics | Semantic. Monitor/Discrimination |
| 94 | 40.5 | -2.9 | -41.4 | ITG | Homolog of node 93 | - |
| 97 | -55.2 | -60.3 | -6.0 | ITG | Semantics and Speech | Film Viewing, Naming (Overt) |
| 98 | 54.2 | -56.9 | -8.6 | ITG | Homolog of node 97 | - |
| 99 | -58.8 | -42.1 | -16.0 | ITG | Orthography and Semantics | Naming (Overt), Reading (Covert), Semantic. Monitor/Discrimination, Word Generation (Overt) |
| 100 | 60.5 | -40.5 | -17.1 | ITG | Homolog of node 99 | - |
| 101 | -54.9 | -30.3 | -27.4 | ITG | Semantics | - |
| 102 | 53.7 | -30.3 | -26.3 | ITG | Homolog of node 101 | - |
| 103 | -32.4 | -16.6 | -32.3 | FuG | Semantics and Speech | Naming (Overt), Semantic. Monitor/Discrimination |
| 104 | 33.1 | -14.6 | -34.1 | FuG | Semantics | Naming (Overt), Semantic. Monitor/Discrimination |
| 105 | -30.6 | -64.4 | -14.1 | FuG | Orthography, Semantics, Speech | Naming (Covert and Overt) |
| 106 | 31.3 | -61.4 | -13.7 | FuG | Language | Naming (Covert and Overt) |
| 107 | -42.3 | -50.9 | -17.3 | FuG | Orthography, Phonology, Semantics, Speech | Naming (Covert and Overt), Phonological. Discrimination, Reading (Covert) and Semantic. Monitor/Discrimination |
| 108 | 42.7 | -49.1 | -18.6 | FuG | Orthography, Semantics | Naming (Covert) |
| 113 | -28.3 | -32.6 | -16.9 | PhG | Semantics | Naming (Overt), Semantic. Monitor/Discrimination |
| 114 | 28.8 | -30.7 | -17.5 | PhG | Homolog of node 113 | Passive Listening, Semantic. Monitor/Discrimination |
| 121 | -54.4 | -39.8 | 4.2 | pSTS | Phonology, Semantics, Speech, Syntax | Passive Listening, Phonological. Discrimination, Reading (Covert), Semantic. Monitor/Discrimination, Word Generation (Covert and Overt) |
| 122 | 52.9 | -36.8 | 3.1 | pSTS | Execution. Speech, Phonology, Semantics, and Speech | Passive Listening, Phonological. Discrimination, Reading (Overt) |
| 123 | -52.4 | -50.3 | 10.8 | pSTS | Orthography, Semantics, Speech, and Syntax | Reading (Covert), Semantic. Monitor/Discrimination |
| 124 | 56.5 | -40.1 | 12.5 | pSTS | Homologues of node 123 | Passive Listening |
| 143 | -46.8 | -64.7 | 25.8 | IPL | Language | Semantic. Monitor/Discrimination, |
| 144 | 53.0 | -54.1 | 24.4 | IPL | Homolog of node 143 | - |
| 155 | -50.4 | -15.8 | 42.1 | PoG | Execution. Speech | Recitation/Repetition. (Overt) |
| 156 | 50.3 | -14.2 | 43.7 | PoG | Execution. Speech | Reading (Overt), Recitation/Repetition. (Overt) |
| 157 | -55.8 | -14.0 | 16.2 | PoG | Execution. Speech | Recitation/Repetition. (Overt) |
| 158 | 55.2 | -10.2 | 15.0 | PoG | Execution. Speech | Recitation/Repetition. (Overt) |

*SFG*, superior frontal gurus; *MFG*, middle frontal gyrus; *IFG*, inferior frontal gyrus; *OrG*, orbital gyrus; *PrG*, precentral gyrus; *STG*, superior temporal gyrus; *MTG*, middle temporal gyrus; *ITG*, inferior temporal gyrus; *FuG*, fusiform gyrus; *PhG*, hippocampal gyrus; *pSTS*, posterior superior temporal sulcus; *IPL*, inferior parietal lobule. *PoG*, postcentral gyrus. The ID is the number of the parcel in the Brainnetome atlas (Fan et al., 2016). Here we only summarized the language-related behavioral domains and paradigm classes; the full behavioral domains and paradigm classes for each node are available at http://atlas.brainnetome.org/bnatlas.php.

Supplementary Table 2 state hubs (> mean nodal degree)

| ID | ID in the Brainnetome Atlas | Anatomy | Module ID | SC nodal degree | Weighted Nodal degree (Edge > 0.2) | | | |
| --- | --- | --- | --- | --- | --- | --- | --- | --- |
|  |  |  |  |  | State 1 | State 2 | State 3 | State 4 |
| 1 | 1 | SFG | 2 | 16 | 4.81 | 13.06 | 4.67 | 3.86 |
| 2 | 2 | SFG | 2 | 12 | 4.21 | 13.62 | 3.55 | 3.83 |
| 3 | 17 | MFG | 2 | 16 | 8.64 | 9.60 | 12.92 | 5.83 |
| 4 | 18 | MFG | 2 | 14 | 8.45 | 11.95 | 10.86 | 5.26 |
| 5 | 29 | IFG | 2 | 15 | 10.60 | 20.29 | 7.51 | 6.58 |
| 6 | 30 | IFG | 2 | 10 | 9.38 | 19.36 | 6.03 | 5.06 |
| 7 | 31 | IFG | 2 | 17 | 10.70 | 19.85 | 8.32 | 6.67 |
| 8 | 32 | IFG | 2 | 9 | 9.49 | 18.07 | 6.02 | 5.30 |
| 9 | 33 | IFG | 1 | 18 | 7.77 | 14.39 | 16.83 | 6.82 |
| 10 | 34 | IFG | 2 | 16 | 6.85 | 18.74 | 15.47 | 6.63 |
| 11 | 35 | IFG | 1 | 22 | 6.52 | 14.20 | 17.60 | 5.93 |
| 12 | 36 | IFG | 2 | 19 | 5.65 | 17.32 | 15.37 | 5.37 |
| 13 | 37 | IFG | 1 | 25 | 6.42 | 17.60 | 7.26 | 5.06 |
| 14 | 38 | IFG | 2 | 21 | 5.55 | 18.95 | 4.72 | 5.02 |
| 15 | 39 | IFG | 2 | 20 | 6.86 | 23.36 | 7.02 | 6.67 |
| 16 | 40 | IFG | 2 | 15 | 7.93 | 23.66 | 7.06 | 6.56 |
| 17 | 53 | PrG | 4 | 15 | 11.50 | 11.49 | 8.26 | 4.17 |
| 18 | 54 | PrG | 4 | 12 | 12.15 | 11.88 | 7.96 | 3.92 |
| 19 | 63 | PrG | 2 | 16 | 8.91 | 22.37 | 6.99 | 6.03 |
| 20 | 64 | PrG | 2 | 13 | 7.75 | 22.90 | 9.17 | 5.41 |
| 21 | 71 | STG | 4 | 17 | 13.58 | 20.11 | 7.51 | 4.80 |
| 22 | 72 | STG | 4 | 16 | 12.03 | 18.78 | 8.10 | 5.08 |
| 23 | 73 | STG | 4 | 22 | 14.96 | 20.40 | 8.36 | 5.42 |
| 24 | 74 | STG | 4 | 21 | 14.02 | 21.38 | 8.20 | 5.90 |
| 25 | 75 | STG | 1 | 17 | 17.63 | 17.38 | 13.99 | 7.31 |
| 26 | 76 | STG | 4 | 16 | 15.80 | 20.14 | 5.79 | 6.57 |
| 27 | 77 | STG | 1 | 27 | 13.50 | 5.39 | 11.00 | 4.43 |
| 28 | 78 | STG | 1 | 23 | 15.19 | 6.45 | 12.49 | 4.77 |
| 29 | 79 | STG | 1 | 21 | 18.17 | 6.48 | 12.82 | 6.31 |
| 30 | 80 | STG | 1 | 15 | 18.59 | 9.73 | 15.02 | 6.37 |
| 31 | 81 | MTG | 1 | 23 | 6.10 | 6.45 | 17.32 | 6.31 |
| 32 | 82 | MTG | 1 | 21 | 5.86 | 5.10 | 15.84 | 5.85 |
| 33 | 83 | MTG | 1 | 22 | 11.20 | 7.34 | 16.04 | 6.42 |
| 34 | 84 | MTG | 1 | 20 | 11.43 | 7.65 | 16.68 | 6.19 |
| 35 | 85 | MTG | 1 | 30 | 6.51 | 15.91 | 12.25 | 3.48 |
| 36 | 86 | MTG | 1 | 23 | 5.11 | 13.76 | 12.03 | 3.32 |
| 37 | 87 | MTG | 1 | 21 | 13.06 | 7.53 | 18.62 | 8.07 |
| 38 | 88 | MTG | 1 | 21 | 15.81 | 7.73 | 19.02 | 7.49 |
| 39 | 89 | ITG | 3 | 13 | 4.12 | 2.80 | 3.76 | 2.73 |
| 40 | 90 | ITG | 3 | 14 | 2.00 | 2.00 | 3.26 | 1.79 |
| 41 | 91 | ITG | 2 | 16 | 8.58 | 11.28 | 5.54 | 4.72 |
| 42 | 92 | ITG | 2 | 14 | 6.71 | 5.72 | 3.90 | 3.12 |
| 43 | 93 | ITG | 1 | 17 | 5.93 | 3.41 | 12.08 | 2.13 |
| 44 | 94 | ITG | 1 | 14 | 5.56 | 2.70 | 9.49 | 2.22 |
| 45 | 97 | ITG | 2 | 27 | 8.47 | 13.91 | 6.31 | 4.97 |
| 46 | 98 | ITG | 2 | 19 | 6.52 | 10.11 | 5.08 | 3.87 |
| 47 | 99 | ITG | 2 | 18 | 8.64 | 5.16 | 11.07 | 5.24 |
| 48 | 100 | ITG | 2 | 18 | 9.38 | 6.89 | 9.15 | 5.35 |
| 49 | 101 | ITG | 3 | 14 | 6.28 | 3.31 | 6.33 | 2.46 |
| 50 | 102 | ITG | 3 | 14 | 5.95 | 4.75 | 2.56 | 2.20 |
| 51 | 103 | FuG | 3 | 27 | 3.27 | 3.68 | 3.46 | 3.21 |
| 52 | 104 | FuG | 3 | 20 | 3.29 | 3.41 | 2.99 | 3.18 |
| 53 | 105 | FuG | 3 | 17 | 9.67 | 1.74 | 5.30 | 1.69 |
| 54 | 106 | FuG | 3 | 14 | 10.48 | 1.93 | 5.41 | 1.67 |
| 55 | 107 | FuG | 3 | 20 | 3.96 | 6.82 | 7.56 | 3.83 |
| 56 | 108 | FuG | 3 | 18 | 5.20 | 2.80 | 6.24 | 2.87 |
| 57 | 113 | PhG | 3 | 10 | 1.93 | 3.72 | 1.90 | 1.88 |
| 58 | 114 | PhG | 3 | 10 | 2.88 | 3.18 | 1.61 | 1.60 |
| 59 | 121 | pSTS | 1 | 16 | 15.75 | 12.75 | 17.13 | 7.54 |
| 60 | 122 | pSTS | 1 | 15 | 15.08 | 11.79 | 16.71 | 6.79 |
| 61 | 123 | pSTS | 1 | 18 | 13.54 | 15.56 | 12.63 | 5.05 |
| 62 | 124 | pSTS | 1 | 15 | 13.97 | 15.78 | 10.87 | 5.42 |
| 63 | 143 | IPL | 1 | 24 | 4.81 | 5.68 | 15.33 | 4.77 |
| 64 | 144 | IPL | 1 | 22 | 5.59 | 4.94 | 15.91 | 4.38 |
| 65 | 155 | PoG | 4 | 16 | 12.65 | 7.31 | 7.64 | 3.64 |
| 66 | 156 | PoG | 4 | 14 | 12.72 | 7.81 | 7.57 | 3.62 |
| 67 | 157 | PoG | 4 | 14 | 14.17 | 18.38 | 9.87 | 5.37 |
| 68 | 158 | PoG | 4 | 15 | 13.42 | 18.91 | 10.02 | 5.80 |
